# The bile acid receptor TGR5 is a modulator of the bone marrow microenvironment

**DOI:** 10.1101/2023.11.22.568250

**Authors:** Alejandro Alonso-Calleja, Alessia Perino, Frédérica Schyrr, Silvia Ferreira Lopes, Vasiliki Delitsikou, Antoine Jalil, Ulrike Kettenberger, Maria Chiara Galavotti, Angela Martins, Charles Bataclan, Dominique P. Pioletti, Kristina Schoonjans, Olaia Naveiras

## Abstract

The gut is an emerging regulator of bone marrow (BM) hematopoiesis, with several molecules involved in this communication. Among them, bile acids (BAs) act as a relay between the microbiota and the rest of the body by activating specific receptors, including Takeda G protein-coupled receptor 5 (TGR5). Although TGR5 has potent regulatory effects in immune cells, its role in the BM remains unclear. Here, we show that TGR5 is expressed across hematopoietic populations, including hematopoietic stem and progenitor cells. While TGR5 deficiency did not affect steady-state hematopoiesis, it mildly impaired cell-autonomous short-term hematopoietic reconstitution. In contrast, TGR5 deletion altered the BM microenvironment. Young male, but not female, *Tgr5*^-/-^ mice showed reduced bone marrow adipose tissue (BMAT) alongside an accumulation of BM adipocyte progenitors, consistent with impaired progenitor differentiation. TGR5 deficiency also blunted BMAT expansion during obesity and aging, with reduced myeloid-lymphoid progenitor ratios in male mice. Together, our results indicate that TGR5 contributes to sex-dependent regulation of BM adiposity and identify it as a potential target for modulating the BM niche.

**Impact statement:** TGR5 loss-of-function reduces regulated bone marrow adipose tissue in male mice, in multiple conditions (young, obese, 1-year old) without affecting that of young females at homeostasis. These data highlight TGR5 as a new player in the bone marrow microenvironment.

## Introduction

The bone, long considered merely a structural organ, is now seen as a highly dynamic tissue in which various processes, including hematopoiesis, bone remodeling, immunity and metabolism are tightly regulated^1–4^. These processes rely on specialized microenvironments, termed niches, that play a critical role in cell fate regulation^5^, with the bone marrow adipose tissue (BMAT) being an emerging niche component^6–10^. The bone marrow (BM) niche regulates hematopoietic stem and progenitor cell (HSPC) quiescence, self-renewal and commitment towards differentiated cells thanks to a combination of soluble factors and cell-cell interactions^1,5,11,12^. Osteoblasts, adipocytes and endothelial cells form the BM stromal niche along with a wide range of BM mesenchymal stromal cell (BMSC)^1,13^. While the involvement of these specific cell types in regulating hematopoiesis is generally accepted, the extent and nature of their action is controversial, especially for cells that form part of the adipocytic lineage. Mature BM adipocytes were first described to have a predominantly negative effect on hematopoietic recovery post-myeloablation^6,14,15^. Subsequent studies showing a positive effect of the BM adipocyte lineage on hematopoietic recovery called these results into question^7,16,17^. Recent work has uncovered a more nuanced role of non-lipidated BM adipogenic precursors in the control of bone homeostasis^8,9^ and in favoring hematopoietic recovery upon irradiation^10,18,19^. As for other adipose depots, BMAT is a plastic tissue and can be remodeled upon injury and dietary or pharmacological stimulation^15,20–24^.

Bone and BM homeostasis are influenced by extramedullary mechanisms, including the incompletely understood gut-BM cross-talk^25–28^. Communication between the gut and distant organs can be mediated by different signaling molecules, including bile acids (BAs), which are metabolites derived from the liver and subsequently modified by the gut microbiome into secondary BAs^29^. These BAs have emerged as powerful signaling molecules that modulate whole-body metabolism^30–32^ through the activation of receptors such as the G protein-coupled receptor Takeda G protein-coupled receptor 5 (TGR5)^30^. BA-TGR5 signaling has been studied in a multitude of organs^31,32^, and profoundly affects adipose tissue physiology through several mechanisms, ranging from beiging^33,34^ and suppression of chronic inflammation in resident macrophages of the white adipose tissue (WAT)^35^, to stimulation of fat oxidation in brown adipose tissue (BAT)^36–38^. In addition, TGR5 expression was observed in human BMSC-derived adipocytes^34^, but the effect of TGR5 on BMAT and on BM function remains unknown. Previous reports have shown that BAs impact fetal and adult hematopoiesis acting as chemical chaperones^39–41^ and, more recently, through direct interaction with myeloid precursors via the vitamin D receptor^42^. However, the effects at the level of TGR5, the most studied membrane BA receptor, remain to be unraveled.

Here, we show that TGR5 is expressed in hematopoietic and non-hematopoietic lineages in the BM. Upon TGR5 deletion, we observed no significant hematopoietic deficit but a mild decrease in short-term, cell-autonomous reconstitution potential after BM transplantation. However, the loss of TGR5 negatively impacted bone microarchitecture and markedly decreased BMAT in young and 1-year-old male mice on a standard chow-diet (CD), as well as in male mice fed a high-fat-diet (HFD). BMAT volume in young female mice at homeostasis was not impacted by TGR5 deletion. Concomitantly, we observed an increase in immunophenotypically-defined adipocyte-progenitor cells (APCs)^15^ coupled to a functional increase in colony-forming-unit fibroblasts and modestly enhanced hematopoietic recovery upon TGR5 loss.

Together, our findings highlight TGR5 as a sex-specific regulator of the BM microenvironment and suggest it as a target for modulating hematopoietic-stromal interactions.

## Results

### TGR5 is expressed in progenitors and differentiated hematopoietic populations

We initially aimed to define the presence of TGR5 in primitive and differentiated hematopoietic populations. For this, we used a GFP reporter mouse model driven by the *Tgr5* promoter (TGR5:GFP)^43^. Flow cytometry analysis of hematopoietic populations showed consistent GFP signal throughout the main hematopoietic stem and progenitor (HSPC) populations in the adult mouse BM, with significant levels of GFP positivity in the lineage negative (Lin^-^), c-Kit^+^, Sca1^+^ cells (LKS), a population highly enriched in murine hematopoietic stem and multipotent progenitor cells (Figure 1A, B).

**Figure 1.**
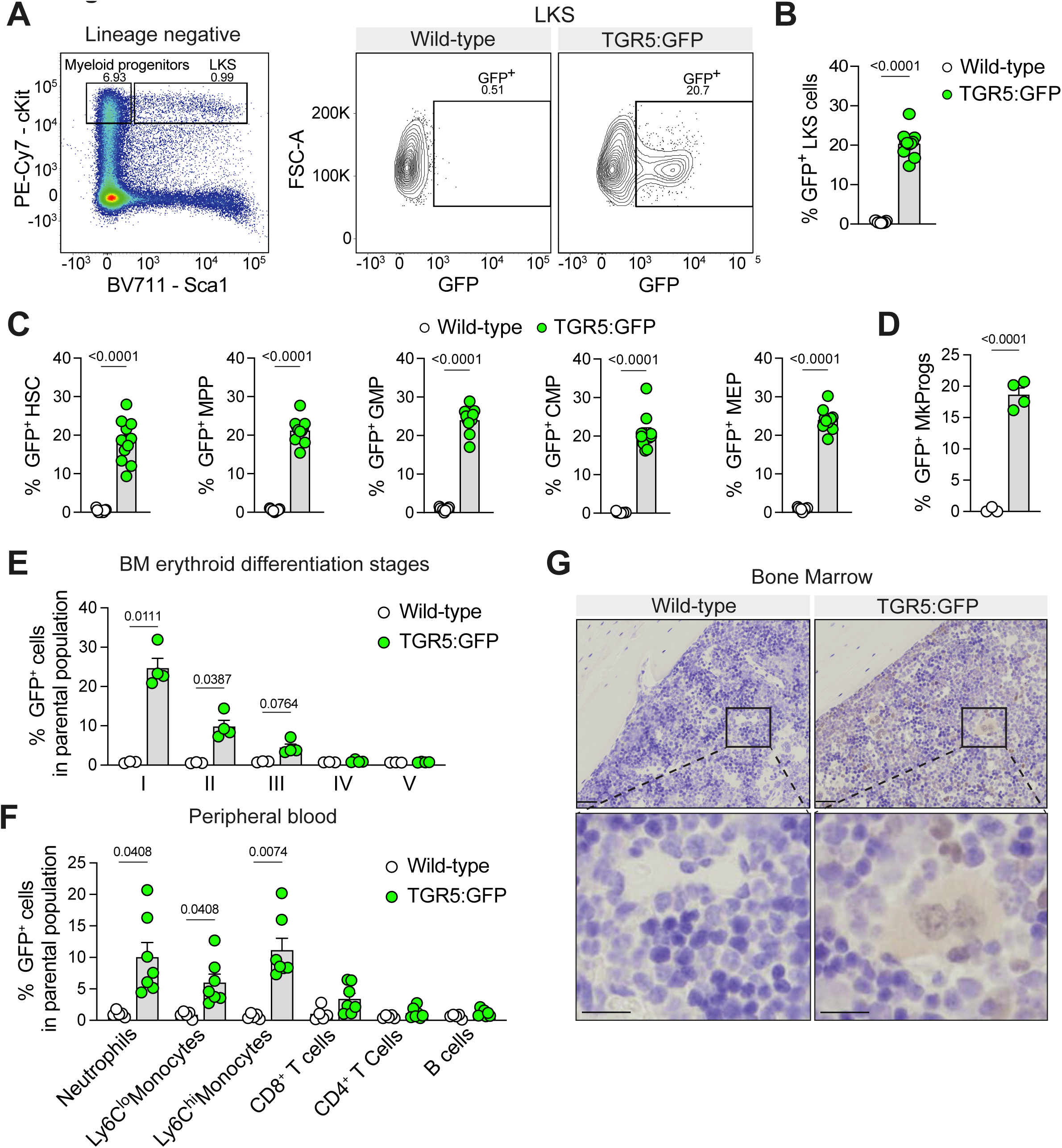
TGR5 is expressed in BM hematopoietic stem and progenitor cells (HSPCs) and in differentiated myeloid populations. (**A**) Representative flow cytometry gating strategy used to identify HSPCs and GFP positivity in TGR5:GFP mice and their wild-type controls. (**B**) Frequency of GFP^+^ cells in the Lineage^-^ cKit^+^Sca1^+^ (LKS) population in the BM of 8- to 12-week-old TGR5:GFP male mice and their controls (n=9 for wild-type mice, n=11 for TGR5:GFP mice). (**C**) Frequency of GFP^+^ cells in hematopoietic stem cells (HSCs: LKS CD150+CD48-), multipotent progenitors (MPPs: LKS CD48+), granulocyte-monocyte progenitors (GMPs: Lineage^-^cKit^+^Sca1^-^CD16/32^+^CD34^+^), common myeloid progenitors (CMPs: Lineage^-^ cKit^+^Sca1^-^CD16/32^low^CD34^+^) and megakaryocyte-erythrocyte progenitors (MEPs: Lineage^-^cKit^+^Sca1^-^CD16/32^-^CD34^low^) in the BM of the mice described in **B**. (**D**) Frequency of GFP^+^ cells in megakaryocyte progenitor (MkProgs) in the BM of the mice described in **B**. (**E**) Frequency of GFP^+^ cells in the different stages of erythropoiesis in the BM of 8- to 12-week-old male TGR5:GFP mice and their wild-type controls (n=3 for wild-type mice, n=4 for TGR5:GFP mice); axis represents % GFP^+^ cells within the parental population. (**F**) Frequency of GFP^+^ cells in peripheral blood cell populations of 8- to 12-week-old male TGR5:GFP mice and their wild-type controls (n=5 for wild-type mice, n=7 for TGR5:GFP mice). (**G**) Immunodetection of GFP^+^ cells by DAB IHC in BM of 8- to 12-week-old TGR5:GFP male mice (n=2 for wild-type mice, n=3 for TGR5:GFP mice). Scale bar: 50 µm in low magnification images, 20 µm in the corresponding digitally zoomed images. Results represent the mean ± s.e.m., n represents biologically independent replicates. Unpaired, two-tailed Student’s t-test (**B**, **C**, **D**), multiple two-tailed Student’s t-test with Holm-Sidak’s multiple comparison correction (**E**, **F**) was used for statistical analysis. P values are indicated.

We detected comparable frequencies of GFP^+^ cells in populations downstream of the LKS gate, namely hematopoietic stem cells (HSC) and multipotent progenitors (MPP) (Figure 1C). Similar frequencies were observed in myeloid progenitors, with no marked differences between the different subpopulations, namely granulocyte-monocyte progenitors (GMP), common myeloid progenitors (CMP) and megakaryocyte-erythrocyte progenitors (MEP) (Figure 1C), as well as in megakaryocyte progenitors (MkProgs) (Figure 1D and Figure 1-figure supplement 1A). Regarding erythropoiesis^44^, we observed a decreasing proportion of GFP-positive cells as cells matured (Figure 1E and Figure 1-figure supplement 1B). Analysis of fully differentiated hematopoietic populations in peripheral blood showed that TGR5:GFP is predominantly expressed in myeloid lineage cells, with lower expression in lymphocytes (Figure 1F and Figure 1-figure supplement 2A). GFP was detected by immunohistochemistry in the BM (Figure 1G), spleen (Figure 1-figure supplement 2B) and, sparsely, in thymus (Figure 1-figure supplement 2C), confirming the expression of TGR5 in different immune organs. Taken together, these findings indicate that TGR5 is present throughout HSPC populations and is mostly restricted to the myeloid lineage upon differentiation.

### TGR5 is not essential for steady-state hematopoiesis nor cell-autonomous hematopoietic stem cell function but modulates short-term BM repopulation

We next hypothesized a role for TGR5 in hematopoietic progenitors and aimed to define the impact of TGR5 loss-of-function in hematopoiesis. Flow cytometry analysis of the BM of young male germline TGR5 knock-out (*Tgr5^-/-^*) mice showed no significant alteration in the percentage (Figure 2A) or absolute number (Figure 2B) of HSPC populations compared to their controls (*Tgr5^+/+^*). Functionally, we could not detect differences in hematopoietic colony-forming unit (CFU) potential, and thus functional blood cell progenitors, between *Tgr5^-/-^* male mice and their *Tgr5^+/+^* counterparts (Figure 2-figure supplement 1A). Although we observed a nearly 20% increase in the average number of total CD45^+^ cells per hindlimb collected from *Tgr5^-/-^* male mice (Figure 2C), this did not result in an increase in CFUs per hindlimb (Figure 2-figure supplement 1B). Complete blood counts (CBC) of peripheral blood showed a modest decrease in eosinophils with no change in other white blood cell populations in *Tgr5^-/-^* male mice (Figure 2D, E). Additionally, we observed a moderate increase in the number of red blood cells, but no change in hemoglobin or platelet levels, in *Tgr5^-/-^* male mice (Figure 2F-H). This differs from recently published work, which showed a modest gain in circulating platelets in *Tgr5^-/-^* mice^45^. Female mice behaved similarly, not showing differences in BM HSPC populations (Figure 2-figure supplement 1C). CBC of young female mice showed no differences in circulating cells between both genotypes, except for monocytes, in which a moderate decrease was observed in the *Tgr5^-/-^* mice (Figure 2-figure supplement 1D-H). Taken together, these results indicate that TGR5 is not required in steady-state hematopoiesis, in neither male nor female young mice.

**Figure 2.**
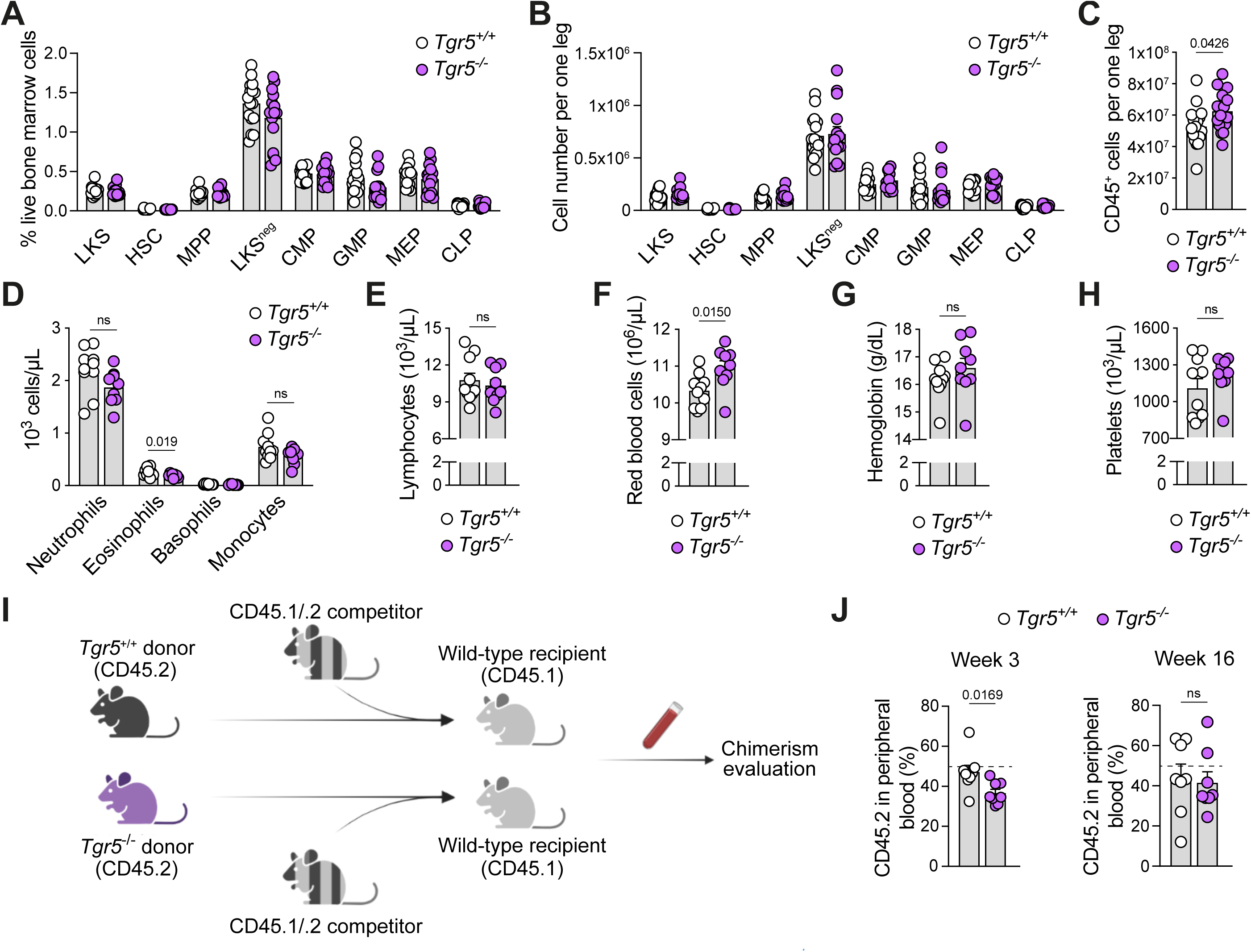
Loss of TGR5 does not impair steady state hematopoiesis but modifies short-term hematopoietic progenitor engraftment. (**A**) Frequency of hematopoietic stem and progenitor (HSPC) populations in the BM of 8- to 12-week-old *Tgr5^+/+^* and *Tgr5*^−/−^ male mice expressed as percentage of cells over total live cells acquired (n=16 for *Tgr5^+/+^* and n=15 for *Tgr5*^−/−^ mice). (**B**) Total number of cells per one leg (hip, femur and tibia) for HSPC populations in the BM of the mice described in **A**. (**C**) Total number of CD45^+^ cells per one leg (hip, femur and tibia) of the mice described in **A**. (**D-H**) Complete blood counts in peripheral blood of 8- to 12-week-old *Tgr5^+/+^* (n=10) and *Tgr5*^−/−^ (n=9) male mice; myeloid cells (**D**), lymphocytes (**E**), red blood cells (**F**), hemoglobin (**G**), and platelets (**H**). (**I**) Workflow depicting the primary competitive transplant setting. (**J**) Flow cytometry analysis of peripheral blood chimerism in primary transplantation recipients (n=8, 8- to 12-week-old *Tgr5^+/+^* and *Tgr5*^−/−^ male donors transplanted into 2-3 CD45.1 recipient mice, treated as technical replicates per donor) at 3 (short-term progenitor readout) and 16 weeks (stem cell contribution readout) post-transplant. Axis represents % CD45.2 donor cells over the sum of CD45.2 donor and CD45.1/.2 competitor events. Dashed line refers to the expected contribution. Results represent the mean ± s.e.m., n represents biologically independent replicates unpaired, two-tailed Student’s t-test (**C**, **D**, **F**, **J**) was used for statistical analysis. P values (exact value) are indicated, ns indicates non-significance.

In the next step, we aimed to define the cell-autonomous effects of a TGR5 germline deletion in hematopoietic stem and progenitor cells in response to stress. To investigate this, we examined the hematopoietic repopulating capacity of *Tgr5^-/-^* and *Tgr5^+/+^* total BM cells when co-transplanted with competitor CD45.1/.2 BM into lethally irradiated wild-type recipients (CD45.1) (Figure 2I). Upon primary transplantation, *Tgr5^-/-^* donor cells showed an average 23% reduction in hematopoietic short-term repopulating capacity as compared to *Tgr5^+/+^* BM cells, the latter approaching the expected 50% average reconstitution capacity in the competitive transplant assay (47% for *Tgr5^+/+^* BM donors reduced to 36% average for *Tgr5^-/-^*, p = 0.0169, Figure 2J, left panel). The decreased short-term hematopoietic repopulating capacity for *Tgr5^-/-^* donor BM cells was absent at later timepoints, when hematopoietic stem cells, not progenitors, are responsible for reconstitution (Figure 2J, right panel). Further, we did not observe any difference in repopulating capacity in the secondary or tertiary transplant in the wild-type donors (Figure 2-figure supplement 1I), with an overall balanced contribution of the lymphoid and myeloid lineage throughout (Figure 2-figure supplement 1J, K), which further indicates preserved hematopoietic stem cell function. Taken together, these findings indicate that *Tgr5^-/-^* BM cells exhibit a mild short-term deficit in repopulating capacity that is corrected over serial transplantation in wild-type recipients, with no detectable long-term impairment of hematopoietic stem cell function.

### TGR5 regulates the BM microenvironment

Based on our findings that functional hematopoietic progenitors are quantitatively conserved at homeostasis and that the cell-autonomous effects on short-term hematopoietic progenitors disappear on serial transplantation (i.e., exposure to a wild-type BM niche), we hypothesized that an altered BM microenvironment in *Tgr5^-/-^* animals may account for this phenotype. In agreement with the literature^46^, we found that young adult *Tgr5^-/-^* male mice display a decrease in bone volume fraction (BV/TV) and trabecular number (Tb.N.) with no changes in trabecular thickness (Tb.Th.) or separation (Tb.Sp.) at the distal femur (Figure 3A,B). In addition, no alterations in cortical parameters were found at the level of the femoral diaphysis (Figure 3-figure supplement 1A, B), as evaluated by micro-computed tomography (µCT). We could not detect any changes in bone microarchitecture in the spine at the level of L4 in young male *Tgr5^-/-^* mice (Figure 3-figure supplement 1C, D). These data suggest that the role of TGR5 on bone microarchitecture diverges based on skeletal location, which is in line with the recently described differential regulation of the axial and appendicular skeletons^47,48^.

**Figure 3.**
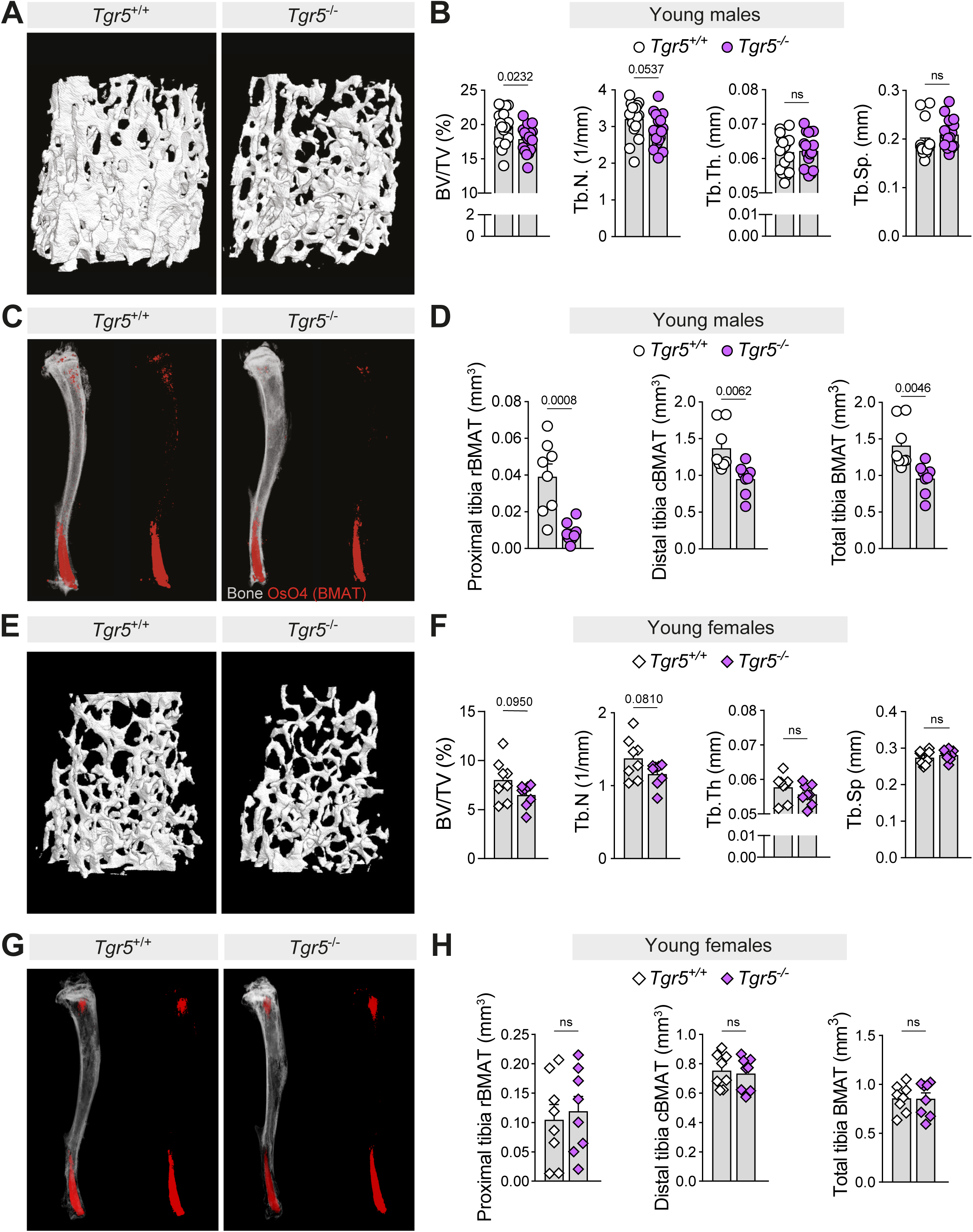
Loss of TGR5 alters the BM microenvironment in male but not in female mice. (**A**) Representative native µCT-derived 3D reconstructions of the trabecular structure in the distal femoral metaphysis of 8- to 12-week-old *Tgr5^+/+^* and *Tgr5*^−/−^ male mice. (**B**) µCT-derived bone morphometry measurements of the trabecular structure in the distal femoral metaphysis of 8- to 12-week-old *Tgr5^+/+^* and *Tgr5*^−/−^ male mice (n=17 for both *Tgr5^+/+^* and *Tgr5*^−/−^ mice); shown are bone volume/total volume (BV/TV), trabecular number (Tb.N.), trabecular thickness (Tb.Th.) and trabecular separation (Tb.Sp.). (**C**) Representative contrast-enhanced µCT-derived 3D reconstructions of osmium tetroxide (OsO_4_)-stained BMAT in the tibiae of 8- to 12-week-old *Tgr5^+/+^* and *Tgr5*^−/−^ male mice. (**D**) Quantification of the OsO_4_-stained bone marrow adipose tissue (BMAT) content in 8- to 12-week-old *Tgr5^+/+^* and *Tgr5*^−/−^ male mice (n=8 for both *Tgr5^+/+^* and *Tgr5*^−/−^ mice). OsO_4_-stained regulated BMAT proximal of tibiofibular junction was classified as “proximal rBMAT”; OsO_4_-stained constitutive BMAT distal of tibiofibular junction was classified as “distal cBMAT”. (**E**) Representative native µCT-derived 3D reconstructions of the trabecular structure in the distal femoral metaphysis of 8-week-old *Tgr5^+/+^* and *Tgr5*^−/−^ female mice. (**F**) µCT-derived bone morphometry measurements of the trabecular structure in the distal femoral metaphysis of 8-week-old *Tgr5^+/+^* and *Tgr5*^−/−^ female mice (n=8 for both *Tgr5^+/+^* and *Tgr5*^−/−^ mice); parameters shown analogously to **B**. (**G**) Representative contrast-enhanced µCT-derived 3D reconstructions of OsO_4_-stained BMAT in the tibiae of 8-week-old *Tgr5^+/+^* and *Tgr5*^−/−^ female mice. (**H**) Quantification of the OsO_4_-stained BMAT content in 8-week-old *Tgr5^+/+^* and *Tgr5*^−/−^ female mice (n=8 for both *Tgr5^+/+^* and *Tgr5*^−/−^ mice). Proximal and distal BMAT were defined as in **D**. Results represent the mean ± s.e.m., n represents biologically independent replicates. Unpaired, two-tailed Student’s t-test **(B, D, F, H)** was used for statistical analysis. P values (exact value) are indicated, ns indicates no statistical significance.

To further study the overall composition of the BM microenvironment, we used osmium tetroxide (OsO_4_) contrast-enhanced µCT^49^ to visualize the tibial BMAT. The BMAT in the proximal tibia, also known as regulated BMAT (rBMAT), is regarded as the most dynamic and stimuli-responsive BMAT in opposition to that found in the distal tibia, which is enriched in the less responsive, constitutive BMAT (cBMAT)^50^. This analysis revealed a profound reduction in proximal tibia rBMAT in young adult *Tgr5^-/-^* male animals, as well as a less marked decrease in distal tibia cBMAT compared to age-matched controls (*Tgr5^+/+^*) (Figure 3C, D). Young *Tgr5^-/-^*females showed a trend towards lower trabecular BV/TV in the appendicular skeleton (Figure 3E, F), a smaller cross-sectional area of the femoral diaphysis (Figure 3-figure supplement 1E-F) and no differences in L4 (Figure 3-figure supplement 1G-H). There was no difference in tibia BMAT between groups (Figure 3G,H), indicating that the effect of TGR5 deletion on BMAT in young mice is sexually dimorphic and is congruent with recent work pointing towards a differential regulation of BMAT in male and female mice^51^.

We next sought to evaluate whether the decrease in rBMAT with a more preserved cBMAT was exclusive to the appendicular skeleton or was also present in axial structures in male mice. For this, we chose the caudal spine, as this segment of the vertebral column contains a gradient where rBMAT gives way to cBMAT in a proximal-to-distal manner^52^. µCT scanning of OsO_4_-stained caudal spines showed this adipocytic gradient, so-called the yellow-to-red transition at the level of the second caudal vertebrae (CA2) in young male *Tgr5^+/+^* mice. Analogously to our findings for the rBMAT in tibia, *Tgr5^-/-^* male mice presented a marked decrease in BMAT at the level of CA2 (Figure 3-figure supplement 2A-C). Histological preparations of the same spine segments from an independent cohort further confirmed our µCT data (Figure 3-figure supplement 2D). Taken together, our data indicate that TGR5 regulates the composition of the BM microenvironment in a skeletal location- and sex-specific manner.

### TGR5-dependent reduction of BMAT accrual and hematopoietic progenitor myeloid bias associated with age and obesity

As mice lose bone and accumulate BMAT with age^52^, we evaluated the hindlimbs of 1-year-old *Tgr5^-/-^* male mice, expecting that the observed bone loss in young males would persist, as shown in previous work^46^, and BMAT expansion would be blunted. Accordingly, 1-year-old *Tgr5^-/-^*male mice showed a decrease in mineralized tissue in the femoral metaphysis (Figure 4A, B) and, to a milder extent, in the femoral diaphysis and L4 vertebral body (Figure 4-figure supplement 1A-D). Similarly, we observed a reduction in tibia BMAT compared to their age-matched controls (Figure 4C, D). As BMAT expansion has been linked to a myeloid bias in BM progenitors^53–55^ and *Tgr5^-/-^*animals fail to expand this tissue upon aging, we hypothesized that the loss of TGR5 would blunt these changes. In agreement with our hypothesis, we found that the CMP/CLP ratio in 1-year-old *Tgr5^-/-^* male mice was significantly lower than that of their wild-type controls (Figure 4E, F).

**Figure 4.**
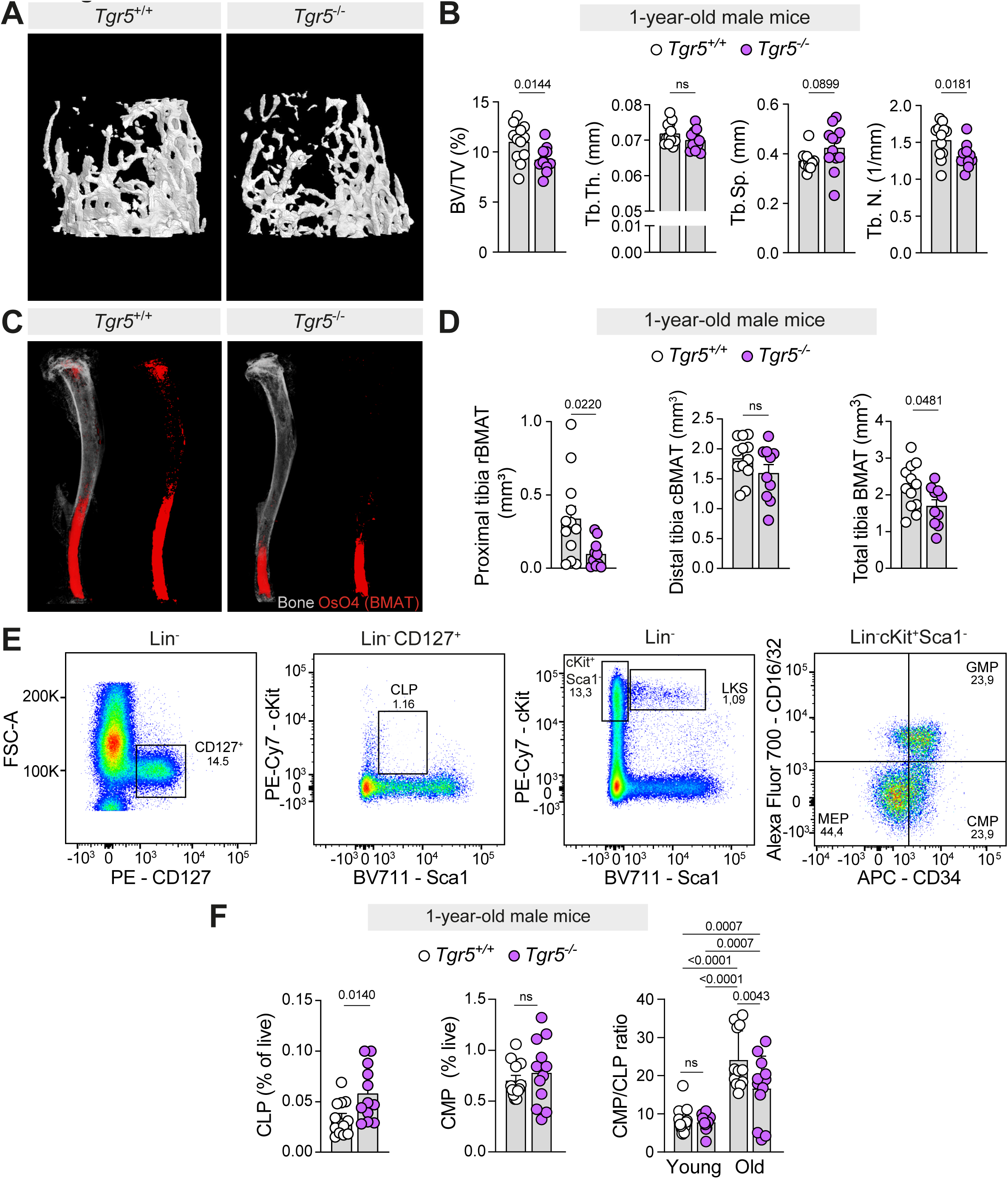
Loss of TGR5 alters BM microenvironment and hematopoietic progenitor myeloid bias in 1-year-old male mice. **(A)** Representative native µCT-derived 3D reconstructions of the trabecular structure in the distal femoral metaphysis of 1-year-old *Tgr5^+/+^* and *Tgr5*^−/−^ male mice. (**B**) µCT-derived bone morphometry measurements of the trabecular structure in the distal femoral metaphysis of 1-year-old *Tgr5^+/+^* and *Tgr5*^−/−^ male mice (n=12 for both *Tgr5^+/+^* and *Tgr5*^−/−^ mice); shown are bone volume/total volume (BV/TV), trabecular thickness (Tb.Th.), trabecular separation (Tb.Sp.) and trabecular number (Tb.N.). (**C**) Representative contrast-enhanced µCT-derived 3D reconstructions of osmium tetroxide (OsO_4_)-stained bone marrow adipose tissue (BMAT) in the tibiae of 1-year-old *Tgr5^+/+^* and *Tgr5*^−/−^ male mice. (**D**) Quantification of the OsO_4_-stained BMAT content 1-year-old *Tgr5^+/+^* and *Tgr5*^−/−^ male mice (n=12 for *Tgr5^+/+^* and n=11 for *Tgr5*^−/−^ mice). OsO_4_-stained regulated BMAT proximal of tibiofibular junction was classified as “proximal rBMAT”; OsO_4_-stained constitutive BMAT distal of tibiofibular junction was classified as “distal cBMAT”. (**E**) Representative gating strategy for common myeloid progenitors (CMP) and common lymphoid progenitors (CLP). (**F**) Frequency of common lymphoid progenitors (CLP), common myeloid progenitors (CMP) and ratio of CMP-to-CLP cells in the BM of 8- to 12-week-old (calculated from the data presented in Figure 2A-B) and 1-year-old *Tgr5^+/+^* and *Tgr5*^−/−^ male mice (n=12 for both *Tgr5^+/+^* and *Tgr5*^−/−^ mice). Results represent the mean ± s.e.m., n represents biologically independent replicates. Unpaired, two-tailed Student’s t-test (**B**, **D**, **F**) and 2-way ANOVA test with Holm-Sidak’s multiple comparison correction (CMP/CLP ratio in **F**) were used for statistical analysis. Exact p-values are indicated; ns indicates non-significance.

Dietary intervention has also been shown to modulate bone microarchitecture and BMAT, with high-fat diet (HFD)-feeding decreasing mineralized tissue and robustly increasing marrow adiposity in general and rBMAT in particular^15,56,57^. To investigate if TGR5 modulates the BM niche upon this dietary intervention, we fed *Tgr5^-/-^* and *Tgr5^+/+^* male mice with an HFD for 12 weeks and evaluated hindlimbs as in 1-year-old animals. Compared to their wild-type controls, *Tgr5^-/-^* animals fed HFD showed a slightly more marked deterioration of their trabecular bone microarchitecture (Figure 5A, B) with minimal impact on cortical parameters (Figure 5-figure supplement 1A, B), and a marked decrease in rBMAT along with a more moderate decrease in cBMAT (Figure 5C, D). Moreover, and in line with data gathered from control chow diet (CD)-fed 1-year-old mice, the CMP/CLP ratio in HFD-fed *Tgr5^-/-^* male mice was similar to that of our younger, CD-fed cohorts (Figure 5-figure supplement 1C).

**Figure 5.**
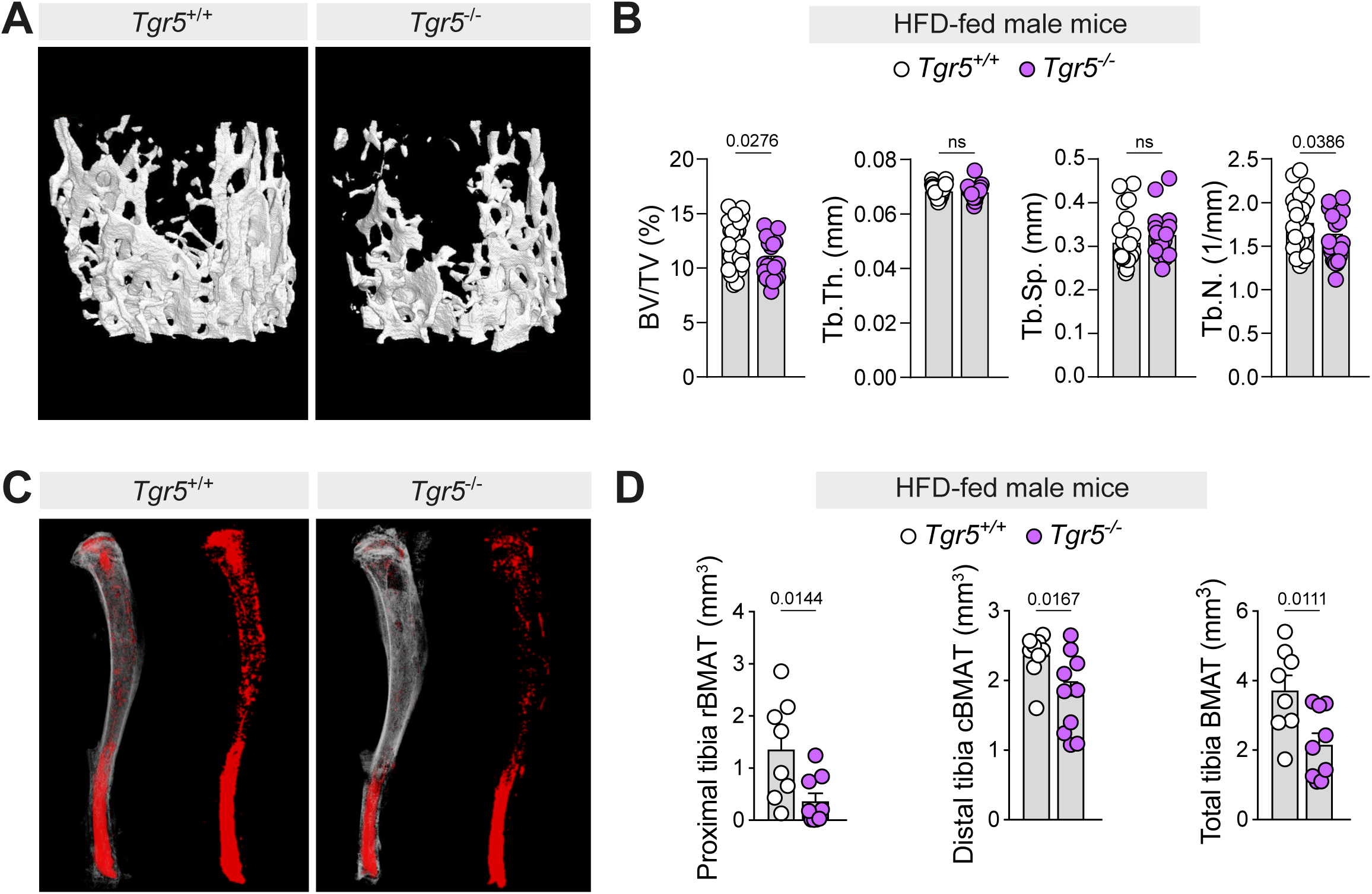
Loss of TGR5 alters the BM microenvironment in HFD-fed male mice. (**A**) Representative native µCT-derived 3D reconstructions of the trabecular structure in the distal femoral metaphysis of *Tgr5^+/+^* and *Tgr5*^−/−^ male mice fed with a high-fat diet (HFD) for 12 weeks. (**B**) µCT-derived bone morphometry measurements of the trabecular structure in the distal femoral metaphysis of HFD-fed *Tgr5^+/+^* and *Tgr5*^−/−^ male mice (20-week-old at the end of the intervention, n=24 for *Tgr5^+/+^* and n=19 for *Tgr5*^−/−^); shown are bone volume/total volume (BV/TV), trabecular thickness (Tb.Th.), trabecular separation (Tb.Sp.) and trabecular number (Tb.N.). (**C**) Representative contrast-enhanced µCT-derived 3D reconstructions of osmium tetroxide (OsO_4_)-stained bone marrow adipose tissue (BMAT) in the tibiae of *Tgr5^+/+^* and *Tgr5*^−/−^ male mice fed with a HFD for 12 weeks. (**D**) Quantification of the OsO_4_-stained BMAT content of mice *Tgr5^+/+^* and *Tgr5*^−/−^ male mice fed with a HFD for 12 weeks (n=9 for *Tgr5^+/+^* and n=10 for *Tgr5*^−/−^ mice). OsO_4_-stained regulated BMAT proximal of tibiofibular junction was classified as “proximal rBMAT”; OsO_4_-stained constitutive BMAT distal of tibiofibular junction was classified as “distal cBMAT”. Results represent the mean ± s.e.m., n represents biologically independent replicates. Unpaired, two-tailed Student’s t-test (**B**, **D**) was used for statistical analysis. Exact p-values are indicated; ns indicates non-significance.

Taken together, our findings indicate a mutual loss of trabecular bone and marrow adiposity in *Tgr5^-/-^* male mice. Specifically, loss of TGR5 decreased trabecular bone in the appendicular skeleton without affecting bone structure at the level of the axial skeleton, except for 1-year-old animals, for which both locations were affected. Conversely, lack of TGR5 markedly reduced BMAT volume in both the appendicular and axial skeleton in male mice. The decrease in BMAT occurred fundamentally at the expense of rBMAT both at homeostasis and under aging- or HFD-induced expansion of BMAT, suggesting that TGR5 might be a key player in the regulation of this tissue.

### Loss of TGR5 promotes adipocyte progenitor accumulation

Given the changes in both BMAT and bone architecture, we hypothesized that BMSCs obtained from *Tgr5^-/-^* mice would present a defect in both adipocytic and osteoblastic differentiation. TGR5 was present in the stroma-enriched CD45^-^Ter119^-^ CD31^-^ population as determined by GFP positivity in the TGR5:GFP reporter model (Figure 6A, B) albeit at lower levels than in HSPCs (Figure 1A-D). Contrary to our hypothesis, *Tgr5^-/-^* BMSCs differentiated more readily into adipocytes, as evidenced by Oil Red O staining (ORO) (Figure 6C, D and Figure 6-figure supplement 1A) and digital holographic microscopy (Figure 6E, F)^58^. Moreover, *Tgr5^+/+^* and *Tgr5^-/-^* BMSCs displayed comparable osteoblastic differentiation, evaluated by alkaline phosphatase staining (ALP) (Figure 6G) and calcium deposition, as evidenced by alizarin red staining (Figure 6H). Taken together, these data indicate that TGR5 deletion does not cause an intrinsic defect in *in vitro* BMSC differentiation.

**Figure 6.**
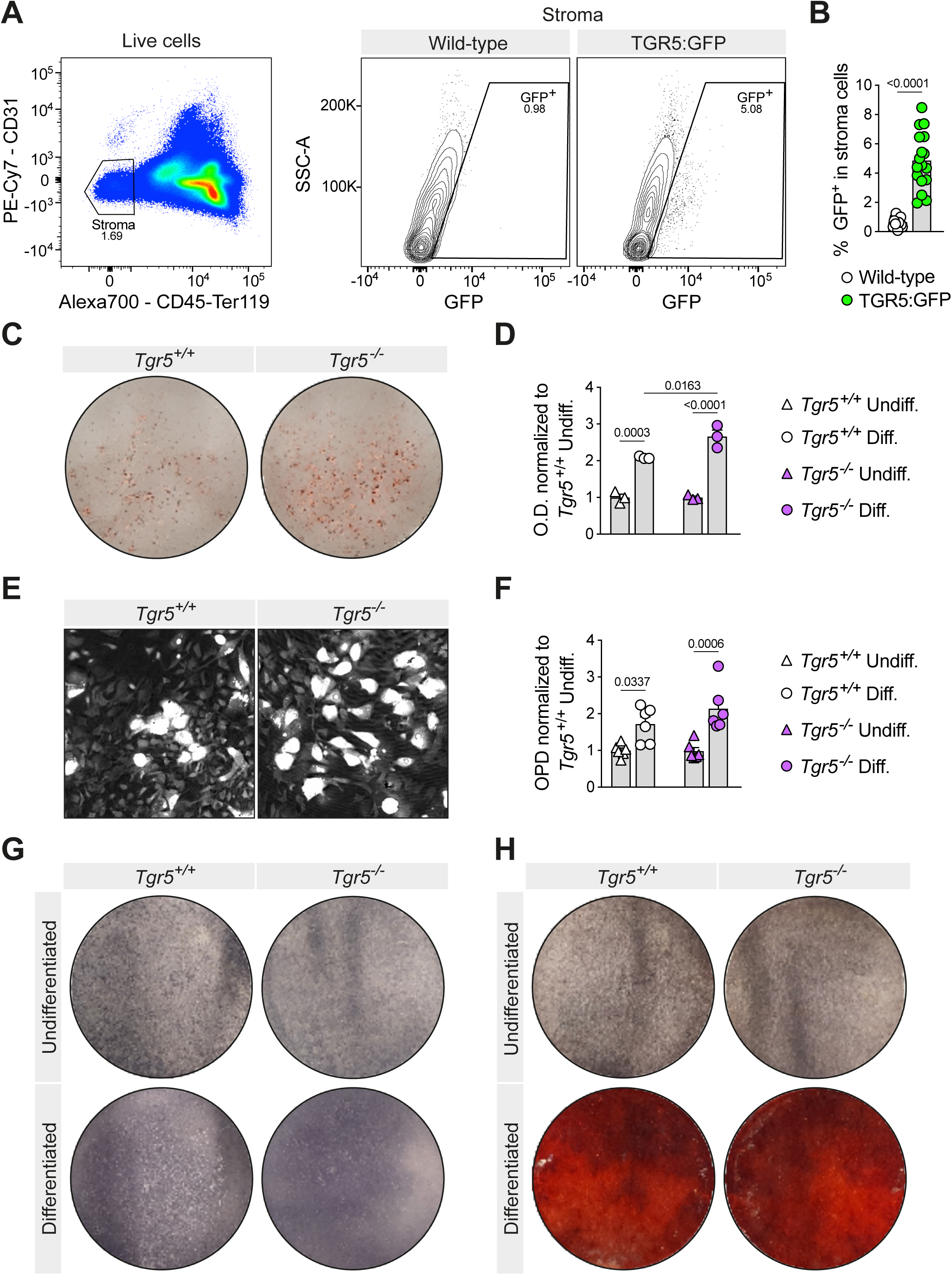
TGR5 is expressed in BM stroma, and its loss does not cause an intrinsic BMSC differentiation defect *in vitro*. (**A**) Representative flow cytometry gating strategy used to identify the stroma-enriched CD45^-^Ter119^-^CD31^-^ gate and the GFP of the cells contained in BM of TGR5:GFP mice. (**B**) Frequency of GFP^+^ cells in the stroma-enriched population of 8- to 12-week-old male TGR5:GFP mice and their wild-type controls (n=15 for wild-type mice, n=17 for TGR5:GFP mice, axis represents % GFP^+^ cells within the parental population). (**C,D**) Representative images (**C**) and spectrophotometric quantification (**D**) of Oil red O (ORO)-stained BMSCs after 7 days of adipocytic differentiation (n=3 for both *Tgr5^+/+^* and *Tgr5*^−/−^ mice). (**E,F**) Digital holographic microscopy imaging of BMSCs after 7 days of adipocytic differentiation showing representative images; lipid inclusions appear as bright white areas in this technique (**E**), and optical path difference (OPD) quantification, indicative of the degree of lipidation (**F**). (**G,H**) Alkaline phosphatase (ALP) (**G**) and alizarin red (AR) (**H**) performed after ALP stains of BMSCs after 7 days of osteoblastic differentiation. All cells were obtained from 8- to 12-week-old *Tgr5^+/+^* and *Tgr5*^−/−^ male mice. Results represent the mean ± s.e.m., n represents biologically independent replicates. Unpaired, two-tailed Students t-test (**B**) and one-way ANOVA with Tukey’s multiple test correction (**D**) or Holm-Šídák’s multiple test correction (**F**) were used for statistical analysis. P values (exact value) are indicated.

Recent reports have revealed that lipodystrophy or specific ablation of pre-adipocytic populations in the BM leads to massive trabecular bone invasion of the medullary cavity, indicating that these populations negatively regulate the mineralized compartment of the BM microenvironment^6,8–10,16,59–61^. We hypothesized that our findings in *Tgr5^-/-^* mice could be explained by an *in vivo* blockage of lipidation for specific adipocyte progenitors, namely APCs, or an increased rate of delipidation of mature adipocytes^19^ that is reversed upon forced induction of *in vitro* adipocytic differentiation. In order to assess this, we analyzed the BM stroma based on the expression of Sca1 and CD24 for APC quantification^15^. In agreement with our hypothesis, we found an increase in the proportion of immunophenotypic APCs (CD45^-^Ter119^-^CD31^-^Sca1^+^CD24^-^) in the stroma of *Tgr5^-/-^* mice (Figure 7A). Finally, we observed an alteration in the clonogenic capacity of BMSCs, evidenced by an increase in fibroblast colony-forming units (CFU-F) in *Tgr5^-/-^* BM compared to wild-type controls (Figure 7B), which agrees with a shift towards less differentiated BM stromal populations in *Tgr5^-/-^* animals. Taken together, our findings indicate that TGR5 loss-of-function perturbs the equilibrium of stromal populations within the marrow cavity and suggest that TGR5 signaling is necessary for BM pre-adipocytes to form and/or retain lipid droplets.

**Figure 7.**
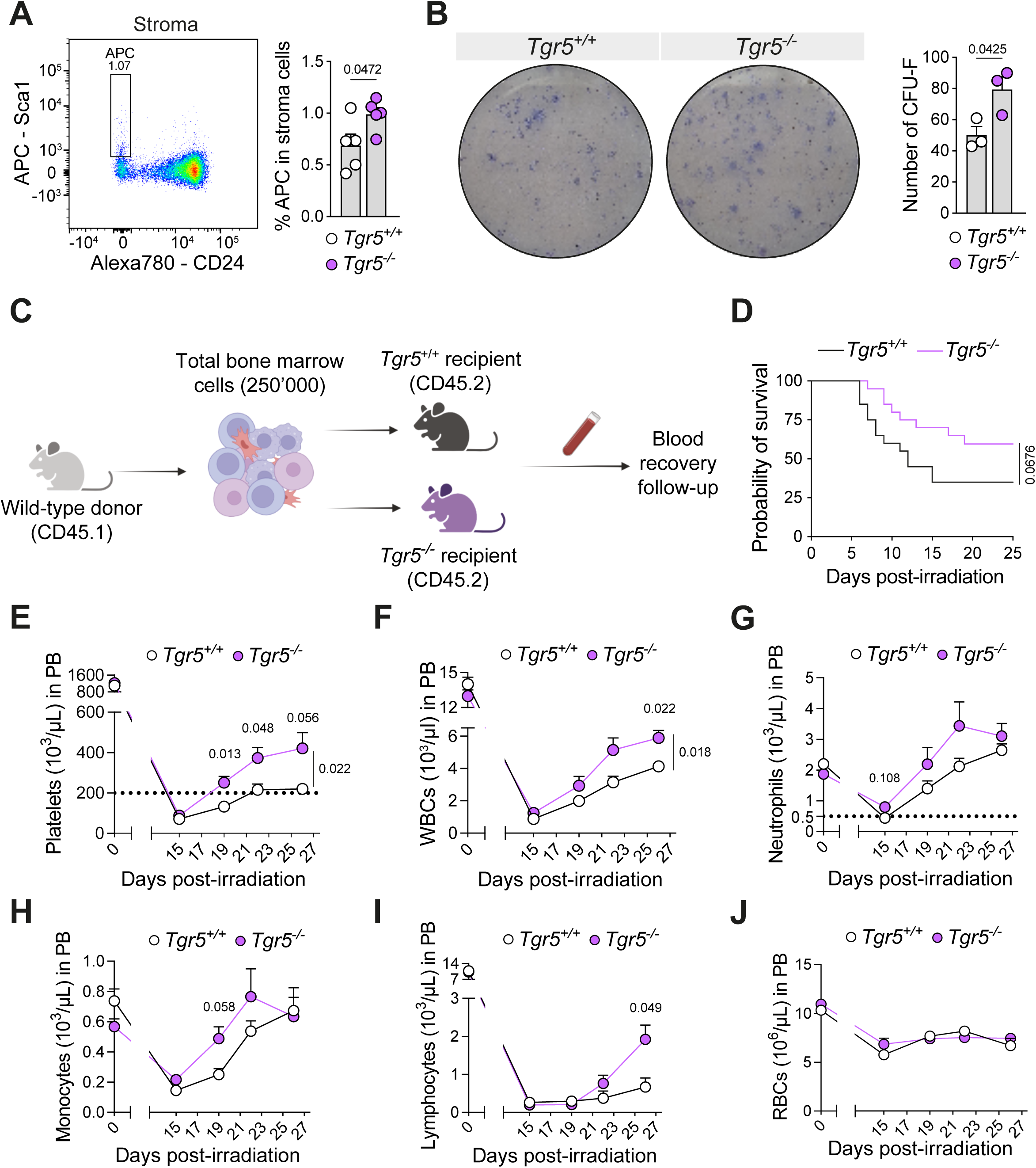
Effect of microenvironmental TGR5 loss in adipocyte progenitor function and hematopoietic recovery. (**A**) Left: representative flow cytometry gating strategy for adipocyte progenitor cells (APC) in the BM stroma gate. Right: percentage of adipocyte progenitor cells (APC) in the CD45^-^Ter119^-^CD31^-^ BM stromal gate (n=5 for both *Tgr5^+/+^* and *Tgr5*^−/−^ mice). (**B**) Left: Representative images for fibroblast colony-forming unit (CFU-F) assay. Right: number of CFU-F obtained from 1 million total BM cells (n=3 both for *Tgr5^+/+^* and *Tgr5*^−/−^ mice). Cells were obtained from 8- to 12-week-old male mice. (**C**) Schematic depiction of the inverse chimera transplantation setup. (**D**) Survival of BM transplant recipients (n=20 for both *Tgr5^+/+^* and *Tgr5*^−/−^ at D0, 8- to 12-week-old male mice in 2 independent experiments). (**E-J**) Peripheral blood recovery evaluated longitudinally by complete blood counts for platelets (**E**), white blood cells (WBC) (**F**), neutrophils (**G**), monocytes (**H**), lymphocytes (**I**) and red blood cells (RBCs) (**J**) (n=7 for *Tgr5^+/+^* and n=12 *Tgr5*^−/−^ 8- to 12-week-old male recipient mice). Results represent the mean ± s.e.m., n represents biologically independent replicates. Unpaired, two-tailed Student’s t-test (**A, B**), Mantel-Cox’s test (**D**) and two-way ANOVA with Holm-Šídák’s multiple comparison correction (**E**-**J**) were used for statistical analysis. For **D**, baseline values (Day 0) were not considered for the analysis of the recovery. P values (exact value) are indicated.

### Effect of microenvironmental TGR5 loss in hematopoietic recovery

Given the enrichment of APCs in the stroma of *Tgr5^-/-^* mice, we next sought to investigate the impact of the associated BM microenvironment alterations in stress hematopoiesis. Lipidated BM adipocytes have been described as negative regulators of hematopoietic recovery^6^, and recent work has shown that non-lipidated adipocytic precursors improve hematopoietic recovery^18^. Considering this, we hypothesized that the shift towards more non-lipidated progenitors in the BM microenvironment of *Tgr5^-/-^*mice would correlate with an improved early hematopoietic recovery upon irradiation and transplantation.

To assess this, we generated inverse chimeras in a lethal irradiation rescue set-up (Figure 7C) and evaluated early hematopoietic recovery using CBCs as a direct readout of the functionality of the hematopoietic system for the first 4 weeks post-irradiation. Our experiments were hampered by challenging microbiological conditions in our facilities, with irradiated mice displaying the *swollen muzzles syndrome*^62^. This led to a mortality that was greater than expected in our cohorts and likely introduced microbiological confounding factors, limiting the interpretation of the results. We reckon these data are nonetheless interesting, as contextualized in the discussion, and present them here to be interpreted with these important caveats into consideration.

We observed a trend towards lower mortality post-transplant in the *Tgr5^-/-^* recipients (Figure 7D, p=0.0679) and faster hematopoietic recovery in *Tgr5^-/-^* recipients upon lethal irradiation and transplantation with wild-type bone marrow donor cells. *Tgr5^-/-^*recipients recovered their platelet levels faster than *Tgr5^+/+^*recipients, showing values consistently above 200.000/µL one week sooner (Figure 7E); platelet levels below this threshold are considered a risk factor for bleeding events^63,64^. Amongst survivors, the recovery was also faster for white blood cells (WBCs) in *Tgr5^-/-^* recipients (Figure 7F) without a clear predominance of any cell type as shown for neutrophils (Figure 7G), monocytes (Figure 7H) or lymphocytes (Figure 7I). We observed no major differences in the recovery of red blood cells (RBCs) (Figure 7J). Taken together, our findings suggest that the loss of TGR5 forms a BM microenvironment that may be beneficial for early hematopoietic recovery in a context of stress hematopoiesis.

## Discussion

In the current study, we aimed to investigate the role of TGR5 on BM homeostasis. In contrast to recent work based on RNA expression data^65^, our flow cytometry analysis using TGR5:GFP reporter mice demonstrated that TGR5 is present in HSPCs, suggesting a more universal role of TGR5 in stem cell homeostasis^66^. Consistent with recently published work reporting TGR5 expression in platelets^45,67^., more differentiated populations, such as megakaryocytes, were also positive for GFP, In contrast, we did not observe an increase in circulating platelets in *Tgr5*^-/-^ mice at baseline, contrary to a previous study^45^, which may be attributable to differences in local animal facility conditions, as it has been shown to impact the microbiome^68,69^. We showed that, in young male mice, TGR5 loss-of-function leads to a relative enrichment of adipocytic precursors in the BM and a simultaneous loss of BMAT, suggesting that TGR5 is necessary for the *in vivo* maturation of BM adipocytes. The latter point was further confirmed by the blunting of the expected BMAT expansion in both 1-year-old and HFD-fed *Tgr5^-/-^* male mice. Further, both these conditions showed a decrease in BM myeloid-lymphoid progenitor ratio, which is in keeping with the relationship of BMAT expansion and myeloid skewing^70^. Young *Tgr5^-/-^* female mice, however, showed no changes in their BMAT, indicating that the effect of TGR5 on BMAT is sexually dimorphic at homeostasis, which is in line with recent reports^51^.

BMAT and the BM adipocyte differentiation axis are gaining attention as a fundamental component of bone and blood physiology, with several recent studies focusing on the BM stroma population as a key player in the maintenance of the balance between bone and BM space^10,71^. Ablation of the BM adipocyte precursor population leads to massive accumulation of bone in the medullary cavity^6,9,16,60^ which in turn reduces the volume available for hematopoietic tissue. In agreement with this model, we observe that *Tgr5^-/-^* male mice display an increase in adipocytic precursors, a decrease in bone, and an expansion of CD45^+^ cells in the BM.

Similar to peripheral adipose depots, BMAT has been described as a highly dynamic tissue that responds to dietary changes, yet has distinct characteristics compared to extramedullary fat, not fully overlapping in its physiology with white or brown adipose tissues^72,73^. For instance, both nutrient deprivation (e.g. caloric restriction and anorexia) and excess (e.g. HFD feeding) result in BMAT expansion^57,74–76^. Although counterintuitive, this nutrient-mediated control of BMAT ensures energy conservation in BM to support the energy-consuming process of hematopoiesis and bone remodeling^77,78^. Another difference between BMAT and peripheral adipose tissues is the absence of HFD-induced pro-inflammatory response in the BMAT compared with the latter^57^. While the role of TGR5 in extramedullary adipose tissues has been extensively studied and shown to be important for beige remodeling^33,34^, lipolysis^34^, and protection against diet-induced immune infiltration of the WAT^32,35^, the effects of an activated TGR5 signaling pathway in BMAT and its impact on hematopoiesis are unknown. Further studies are therefore warranted to define where and how TGR5 acts along the BM adipogenic axis to modulate BMAT, and how these changes modulate hematopoiesis.

We found that young *Tgr5^-/-^* male mice receiving BM transplantation after lethal irradiation recovered faster than their *Tgr5^+/+^* counterparts in a setting of stress hematopoiesis. As stated above, it is important to note that these experiments were carried out in a period during which our irradiated animals suffered from *swollen muzzles syndrome*, associating higher mortality than expected in the early post-transplant period, before hematopoietic toxicity is expected^62^. Therefore, our results should be interpreted within this specific context. While this condition-associated to bacterial colonisation-is a significant limitation of the study, we postulate that it may also carry clinical relevance, as most transplant patients experience at least one episode of febrile neutropenia, often due to a severe infection arising from intestinal bacterial translocation or catheter entry sites^79,80^. While further studies are warranted to clarify how TGR5 influences transplant efficiency in infection-free settings, our findings are conclusive regarding a role for this receptor in the regulation of BMAT. Previous work has shown that *Tgr5* transcript levels increase during *in vitro* differentiation of human BMSCs to adipocytes^34,81^, and BA–mediated TGR5 activation has more recently been shown to suppress BMSC senescence and direct stromal lineage commitment, supporting a broader role for this receptor in BM stromal homeostasis^82^. BMAT adipocytes can also revert to a non-lipidated precursor state^19^. This plasticity may be relevant to the interpretation of our findings, as the BMAT phenotype observed in *Tgr5^-/-^* male mice could result from an accumulation of hematopoietic-supportive adipocyte precursors^18^, a decrease in hematopoietic-quiescence inducing lipidated adipocytes^6^, or a combination of both.

In addition to the role of TGR5, prior studies have linked BAs and transplantation outcomes. Of note, the BA ursodeoxycholic acid (UDCA) is routinely used in clinics in the context of BM transplantation, where supplementation with UDCA has been shown to reduce post-transplant complications and improve overall survival^83^. Available reports have attributed these benefits to the unique feature of the tauro-conjugated form of UDCA (TUDCA) as a chemical chaperone, improving protein folding in fetal liver HSCs^40^ or after acute BM insult^41^, and modulating intestinal antigen presentation to minimize graft-versus-host disease^84^. In addition, UDCA and TUDCA are hydrophilic BAs, and part of their beneficial effects could be a consequence of changes in the hydrophobicity of the BA pool^85,86^. BAs signal through dedicated BA receptors, but the receptor-mediated signaling of UDCA and TUDCA remains yet unclear. Although some studies proposed UDCA as a TGR5 agonist^87–90^, effects occur mainly at supra-physiological doses of UDCA. In addition, the EC_50_ of UDCA for TGR5 is more than 60 times higher than that of the strongest endogenous TGR5 agonist, lithocholic acid (LCA)^91^, indicating that UDCA is a poor TGR5 agonist. Based on this notion, it will be interesting to investigate whether, in this context, the benefits of UDCA upon stress hematopoiesis result from the inability to activate TGR5 signaling in the BMAT.

Novel therapeutic strategies could be envisioned in the future to pharmacologically mimic TGR5 inhibition in the BMAT. Although several selective TGR5 agonists have been developed, only a few TGR5 antagonists have been discovered, namely triamterene^92^, SBI-115^93^ and SBI-364^94^. However, these compounds have only been used *in vitro,* and additional *in vivo* research and optimization will be required to, transiently, pharmacologically block TGR5 in the BM upon transplantation or other conditions linked to stress hematopoiesis, in which a fast production of mature blood cells is required. Our results indicate that, although the exact mechanism remains to be clarified, deletion of TGR5 modulates the BM adipocyte lineage and suggest that strategies aimed at blocking the activity of this receptor might be therapeutically attractive.

## Supporting information

Supplementary Table 1 _ List of Antibodies

Key Resources Table

## Author contribution

AAC, AP and FS: Data curation, Formal analysis, Investigation, Methodology, Writing – original draft, writing – review and editing. SFL, VD, AJ, UK, MCG, AM, CB, DPP: Methodology. KS and ON Conceptualization, Resources, Supervision, Funding acquisition, Validation, Investigation, Writing – original draft, Project administration, Writing – review and editing.

## Acknowledgements

We wish to thank the Histology Core Facility, Center for Phenogenomics, Bioscreening Facilities, Electron Microscopy facilities and BioImaging and Optics Platform at EPFL as well as the In Vivo and Cellular Imaging Facilities at AGORA (UNIL/EPFL) and the Flow Cytometry Facility at AGORA (UNIL/EPFL) for their continuous support and guidance for this project, as well as Dr. Frédéric Schütz (UNIL) for help on statistical analysis. This work was funded by the Ecole Polytechnique Fédérale de Lausanne (EPFL), the University of Lausanne (UNIL), the Swiss National Science Foundation (SNSF N° 310030_189178, Sinergia CRSII5_180317/1 to K.S. and SNSF N° PP00P3_144857, 183725 and 320030-231787 to O.N.), and La Caixa Foundation (to K.S.). Alessia Perino was supported by a fellowship from AXA Research Fund. Frédérica Schyrr was funded by SNSF MD-PhD grant 183986.

## Materials and methods

### Mouse studies and ethical approval

Experiments were carried out in accordance with the Swiss law and with approval of the cantonal authorities (Service Vétérinaire de l’Etat de Vaud, authorization nos. 2990, 2990.1, 2990.2 and 3740).

#### Generation of mouse models and tissue collection

To evaluate the expression of TGR5, we used a transgenic mouse reporter model (TGR5:GFP), that co-expresses human TGR5 and the enhanced green fluorescent protein (GFP) linked by the foot-and-mouth disease virus (*F2a*) cleaving peptide sequence under the control of the endogenous *Tgr5* mouse promoter^43,95^. Mice in these groups were bred in-house at the EPFL facility as offspring of homozygous parents (not littermates) and age-matched as close as possible.

Mice were housed with ad libitum access to water and food and kept under a 12-h dark/12-h light cycle with a temperature of 22 °C ± 1 °C and a humidity of 60% ± 20%. 8- to 12-week-old male mice fed chow diet (CD, SAFE 150) were used for all experiments. 7- to 9-week-old female and 1-year-old male mice fed, both fed CD (SAFE 150) were used where indicated. For the HFD experiment, 8-week-old male mice were fed HFD (Research Diets D12492) for 12 weeks. The whole-body TGR5 knock-out mouse model (*Tgr5^-/-^*) has been previously described^96^. Mice in these groups were bred in-house at the EPFL facility as offspring of homozygous parents (not littermates) and age-matched as close as possible.

C57BL/6J (CD45.2) and C57BL/6J Ly5.1 (CD45.1) mice were bred in-house at the EPFL facility. Double congenic mice (CD45.1/.2) were bred in-house, by F1s from the crossing of CD45.2 and CD45.1 mice.

#### BM transplantation

For transplantation assays, recipient mice were irradiated (lethal x-ray irradiation 8.5 Gy, split in two 4.25 Gy doses 4 h apart using an RS-2000 X-ray irradiator (RAD SOURCE). Transplanted cells (total BM), obtained by crushing the hindlimb bones in a mortar and pestle, were administered the day following irradiation via tail-vein injection. For two weeks after lethal irradiation, mice were provided with paracetamol in drinking water (500 mg Dafalgan in 250 ml water). No antibiotics were provided in drinking water. For the primary competitive transplant setting, 300.000 total BM cells of each the CD45.2 donor (*Tgr5^-/-^* or *Tgr5^+/+^*) and the CD45.1/.2 competitor were co-administered. For the secondary and tertiary transplant, 4 million cells were collected from the primary donor and transplanted into the recipient. For the inverse chimera experiments where the CD45.2 mice were the recipients (*Tgr5^-/-^* or *Tgr5^+/+^*), 250.000 CD45.1 total BM cells were administered.

Blood collection for the experiments described in this work was performed at noon. Blood was obtained from the tail vein by means of a small incision with a scalpel blade and collected into EDTA-coated tubes. Complete blood counts were obtained with an Element HT5 hematology analyzer (Heska, USA).

#### Data reporting

Mice were randomly assigned to control and intervention groups. No statistical methods were used to predetermine sample size but group size was chosen based on variance of previous experiments. The investigators were blinded to genotype allocation during experiments and outcome assessment unless otherwise stated.

#### µCT analysis

To evaluate bone microarchitecture and osmium tetroxide (OsO_4_) stains, a SkyScanner 1276 (Bruker, Belgium) was used. For both applications, a 0.25 mm Al filter was used with a voltage of 200 kV and a current of 55 mA. Samples were wrapped in PBS-soaked paper towels and scanned inside a plastic drinking straw sealed on both ends to avoid drying. Voxel size for both applications was set at 10x10x10 µm^3^.

Bone microarchitecture was evaluated according to the ASBMR guidelines^97^ using a custom CTan (Bruker, Belgium) script for automatic segmentation of trabecular bone in the distal femoral VOI, which was set 100 slices proximal to the distal growth plate and extended 200 slices towards the femoral diaphysis (slice thickness of 0.010 mm). The threshold used to binarize the calcified trabecular issue was 40 on a 0-255 scale. Reconstruction of the scans was performed using NRecon (Bruker, Belgium) and further analysis was performed using CTan (Bruker, Belgium) with the minimum for CS to image conversion set at 0 and maximum set at 0.14. For the analysis of cortical parameters, the midpoint of the femur was determined, and the VOI was defined as the bone 50 slices (slice thickness of 0.010 mm) distal and proximal of the slice corresponding to the midpoint of the bone. All other reconstruction parameters were kept the same as for the analysis of trabecular bone. The threshold used to binarize the calcified trabecular issue was 110 on a 0-255 scale. We provide a representative log file for bones analyzed with the trabecular and the cortical segmentation scripts for reference.

OsO_4_ staining was performed on formalin-fixed, EDTA-decalcified tibiae and caudal spines. For this, bones were kept in 10% formalin for 24 h at 4 °C. Following fixation, the bones were decalcified in PBS-0.5 M EDTA for 2 weeks, refreshing the decalcifying solution on day 7. OsO_4_ staining was performed as previously described^49^. Reconstruction of the scans was performed as for calcified bone except for the minimum and maximum CS range, which was set to 0 and 0.30, respectively. OsO_4_ was segmented using a threshold of 50 on a 0-255 scale. Images and quantification of BMAT were analyzed using Dragonfly software (ORS, Canada).

#### Flow cytometry

All antibodies used for this work as well as their dilutions are listed in the key resources table. All dilutions and cell suspensions were prepared in FACS buffer (2% FBS + 1 mM EDTA in PBS).

#### Histology

Spleens and thymi were fixed in 10% formalin for 24 h at 4 °C. EDTA-decalcified bones (as described previously for OsO_4_ stain), spleens and thymi were cut longitudinally in 3-4 µm sections for staining. For immunohistochemistry (IHC), detection of rabbit anti-GFP was performed manually. After quenching with 3% H_2_O_2_ in PBS 1x for 10 min, a heat pretreatment using 0.1 M Tri-Na citrate pH 6 was applied at 60 °C in a water bath overnight. Primary antibodies were incubated overnight at 4 °C. After incubation of ImmPRESS HRP (Ready to use, Vector Laboratories), revelation was performed with DAB (3,3′-Diaminobenzidine, D5905, Sigma-Aldrich). Sections were counterstained with Harris hematoxylin and permanently mounted. All slides were imaged on an automated slide scanner (Olympus VS120-SL) at 20x or 40x.

### Cell culture

#### Cell extraction

Culture of primary BMSCs was performed as reported in the literature^98^. 8- to 12-week-old male mice were used for these experiments. To isolate BMSCs from the BM, we collected femora and tibiae, cut the epiphyses, and placed the bones in a 200 μL pipette tip with its end cut to fit in a 1.5 mL microcentrifuge tube. Bones were spun at 10.000 g for 10 sec to flush out the BM from the medullary cavity. The BM plug was then transferred to a 15 mL conical tube containing 10 mL of ascorbic-acid free αMEM (containing 10% FBS and 1% penicillin-streptomycin) and this suspension filtered through a 70 μm mesh filter prior to plating in a 100 mm cell culture dish (Corning, USA). Cells were replated 72 h later at a density of 62.500 cells/cm^2^ in tissue-culture treated 12-well plates (i.e., 250.000 cells/well) and grown to confluency prior to differentiation, which usually occurred at one week post-replating. Culture medium was refreshed every other day until confluency^98^.

For CFU-F assays, BM was obtained in a similar manner and plated at a density of 1 million cells per well of a tissue-culture treated 6-well plate (i.e., 100.000 cells/cm^2^). CFU-F cultures were kept in αMEM containing 10% FBS and 1% penicillin-streptomycin and stained on day 7 post-plating with 0.1% toluidine blue in 10% formalin.

#### Media composition and differentiation cocktail

Differentiation into osteoblasts was started once confluency was reached. For osteoblastic differentiation, the base culture medium of ascorbic-acid free αMEM containing 10% FBS and 1% penicillin-streptomycin was supplemented with L-ascorbic acid (final concentration 8 mM) and β-glycerophosphate (final concentration 1 mM) and refreshed every other day until the endpoint of the assay. Dexamethasone was not included in the differentiation cocktail, as per the published protocol we followed^98^.

Differentiation into adipocytes was started once confluency was reached. For adipocytic differentiation, base culture medium was high-glucose (4.5 g/L) DMEM supplemented with 10% FBS and 1% penicillin-streptomycin. Adipocytic induction medium containing dexamethasone (1 μM), isobutylmethylxanthine (IBMX, 0.5 mM), insulin (2 μM) and rosiglitazone (20 μM) was prepared. Cells were cultured in adipocytic differentiation medium for 4 days with refreshing on day 2. On day 4, culture medium was changed to maintenance medium, prepared by supplementing the base medium with only insulin (2 μM) and rosiglitazone (20 μM). Cells were kept in maintenance medium until the endpoint (day 7 post-differentiation) with no medium refreshing.

#### Differentiation stains

For alkaline phosphatase (ALP) stain of osteoblast cultures, one tablet of SigmaFast (Sigma-Aldrich) was dissolved in 10 mL distilled water. Osteoblasts were fixed in 10% formalin for 1 min, washed once with PBS-0.05% Tween 20 and incubated in the staining solution for 6 min. Cells were then washed with PBS-0.05% Tween 20 and imaged.

For alizarin red (AR) stain of calcium deposits in osteoblastic cultures after ALP staining, cells were washed once with distilled water and incubated for 45 min with AR stain (prepared fresh with 2 g of AR in 100 mL distilled water and pH adjusted to 4.2). Cells were then washed with distilled water 4 times and imaged.

For Oil Red O (ORO) staining of adipocyte cultures, we fixed the cells in 10% formalin for 1 h at room temperature. In the meantime, we filtered the ORO stock solution through filter paper as described in the literature^98^. Cells were stained with ORO for 15 min, washed 6 times with distilled water, imaged, and incubated for 10 min with 100% isopropanol to solubilize the stain. ORO was quantified using an iD3 plate reader (Agilent, USA) set to measure absorbance at 490 nm^98^.

### Digital holographic microscopy

Digital holographic imaging was performed in black wall 96-well imaging plates (Corning, 353962) using Transmission DHM® T1000 (Lyncée Tec, Switzerland). Plates were pre-coated with Poly-ornithine (Sigma,100 mg/L) to prevent cell detachment. The cells were imaged live and without prior liquid manipulation using a 20x/0.4 NA microscope objective. The best-focus phase images were reconstructed automatically in MATLAB (MathWorks, USA) from the acquired holograms and the average quantitative phase signal or optical path difference (OPD) was automatically measured using a fixed threshold value.

### Statistics

Differences in groups were assessed as stated in the legend for each figure. GraphPad Prism 9 was used for all statistical analyses. Before data analysis, outliers were checked, identified and excluded by using the Grubbs method (α value = 0.05) in GraphPad Prism 9. The data sheets provided with this work include the points flagged as outliers.

In Figure 5B and C, one animal from each genotype was excluded from analysis in “proximal tibia BMAT” and “total tibia BMAT” as the proximal epiphyses were avulsed upon dissection; these animals were kept for “distal BMAT” analyses as this part of the bone was intact.

The studies were either replicated and/or the results were confirmed by using different approaches (flow cytometry and IHC, for example) yielding similar results. All P values ≤ 0.05 were considered significant. When the exact value is not provided, *P ≤ 0.05, **P ≤ 0.01, ***P ≤ 0.001 and ****P ≤ 0.0001.

## Figure legends – figure supplements

**Figure 1-figure supplement 1.**
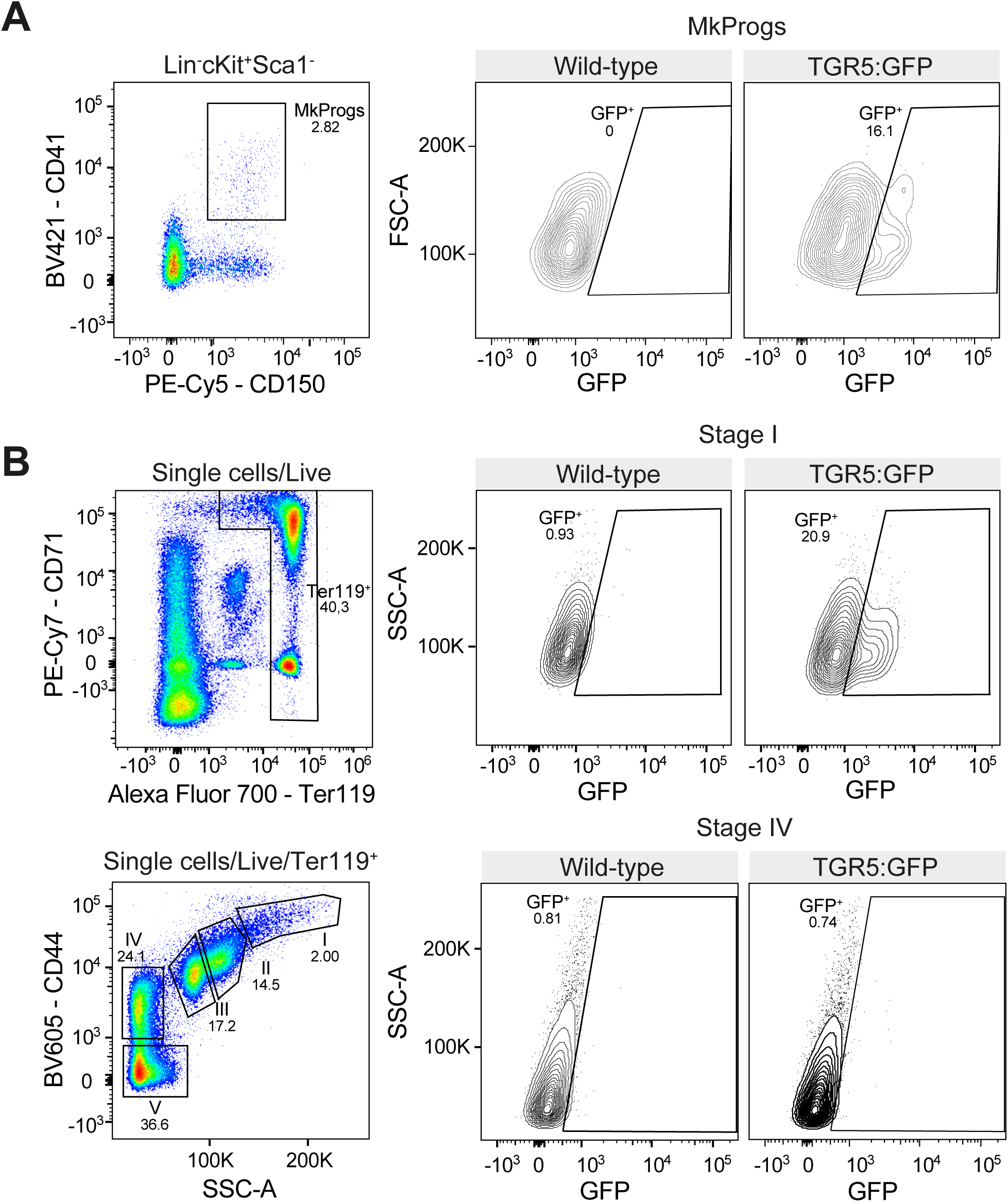
TGR5 is expressed in hematopoietic progenitors in the bone marrow. (**A**) Representative flow cytometry gating strategy used to identify megakaryocyte progenitors (MkProgs) in BM and their GFP positivity in TGR5:GFP mice. (**B**) Representative flow cytometry gating strategy used to identify cells in the different stages of erythropoiesis in BM and their GFP positivity in TGR5:GFP mice.

**Figure 1-figure supplement 2.**
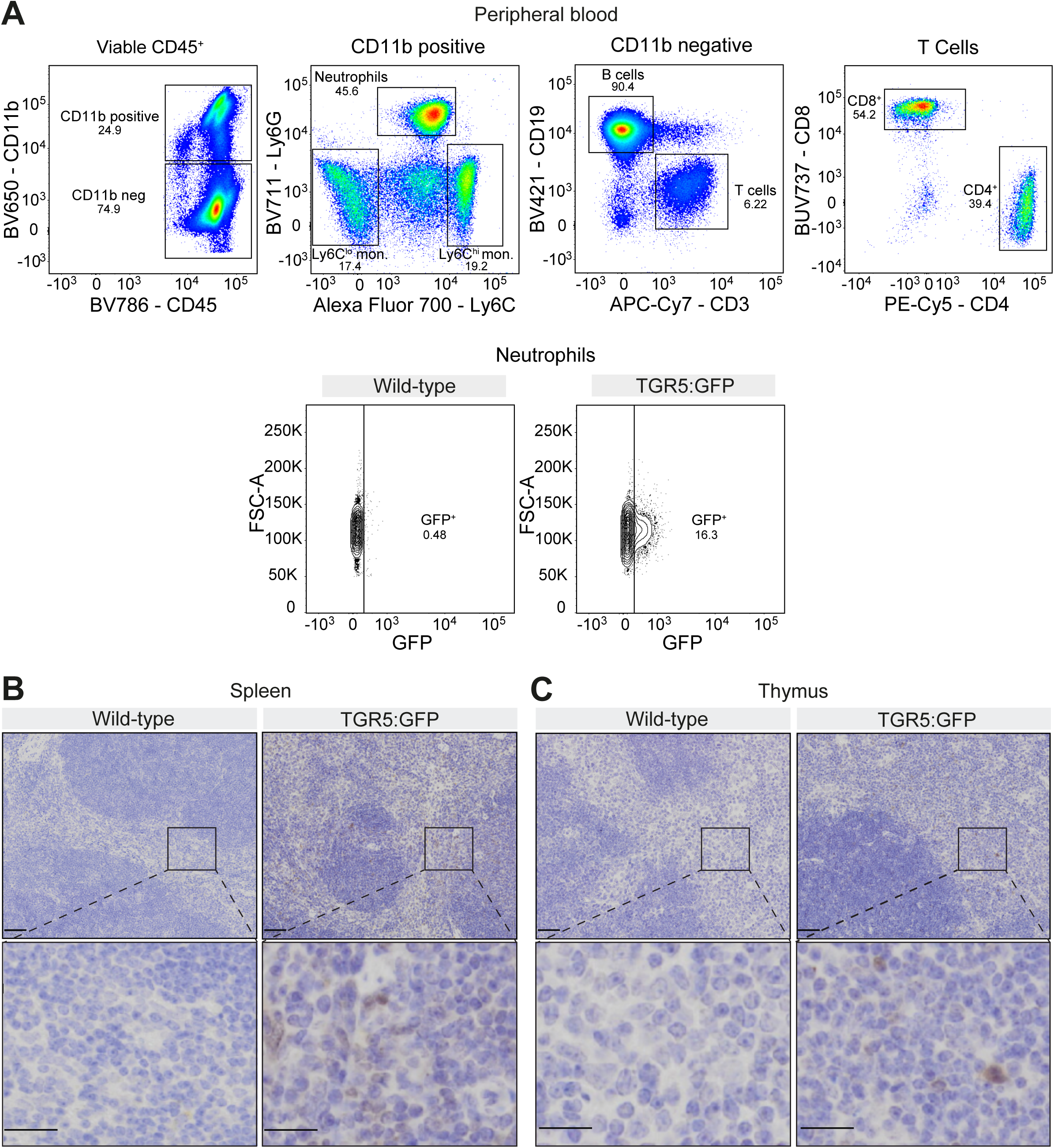
TGR5 is expressed in spleen, thymus and differentiated hematopoietic populations. (**A**) Representative flow cytometry gating strategy used to identify differentiated populations in peripheral blood and GFP positivity in TGR5:GFP mice. (**B**, **C**) Immunodetection of GFP^+^ cells by DAB IHC in spleen (**B**) and thymus (**C**) of 8- to 12-week-old male TGR5:GFP mice (n=2 for wild-type mice, n=3 for TGR5:GFP mice). Scale bar: 50 µm in low magnification images, 20 µm in the corresponding digitally zoomed images.

**Figure 2-figure supplement 1.**
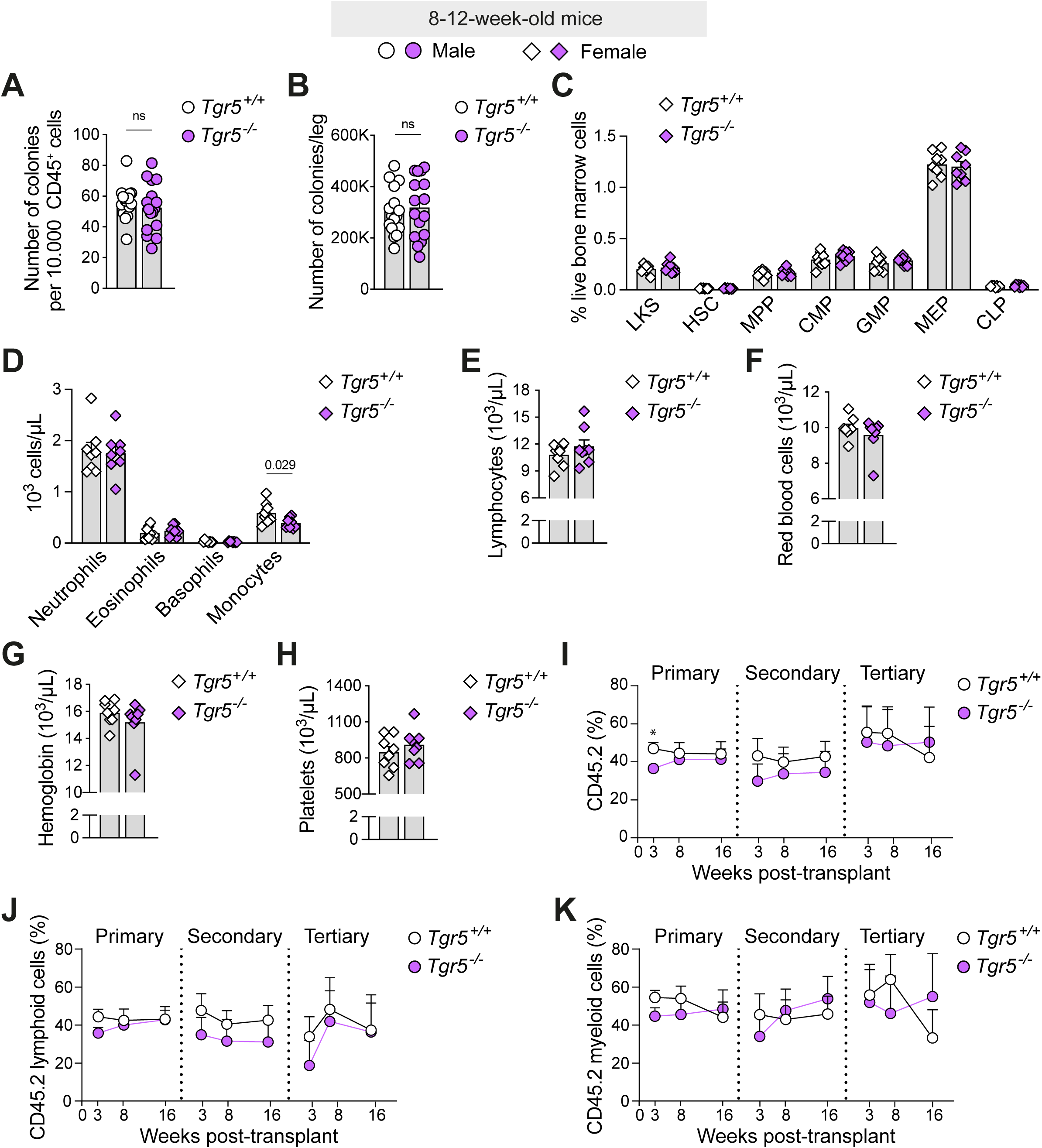
Loss of TGR5 does not impair steady state hematopoiesis but modifies short-term hematopoietic progenitor engraftment. **(A, B)** Number of colonies per 10.000 BM CD45^+^ cells (A) and per leg (B) (n=16 for both *Tgr5^+/+^* and *Tgr5*^−/−^ mice). **(C)** Frequency of hematopoietic stem and progenitor (HSPC) populations in the BM of 8-week-old *Tgr5^+/+^* and *Tgr5*^−/−^ female mice expressed as percentage of cells over total live cells acquired (n=8 for both *Tgr5^+/+^* and *Tgr5*^−/−^ mice). (D-H) Complete blood counts in peripheral blood of 8-week-old *Tgr5^+/+^* and *Tgr5*^−/−^ female mice; myeloid cells (D), lymphocytes (E), red blood cells (F), hemoglobin (G), and platelets (H). (I-K) Flow cytometry analysis of peripheral blood chimerism expressed as the percentage of CD45.2 (I), CD45.2 lymphoid (**J**) and CD45.2 myeloid (**K**) cells of serial BM transplantation experiments (n=8, 8-12-week-old *Tgr5^+/+^* and *Tgr5*^−/−^ male donors transplanted into 3 CD45.1 recipient mice each) at 3, 8 and 16 weeks post-transplant. Results represent the mean ± s.e.m., n represents biologically independent replicates. Unpaired, two-tailed Student’s t-test was used for statistical analysis. *, p<0.05, ns indicates non-significance.

**Figure 3-figure supplement 1.**
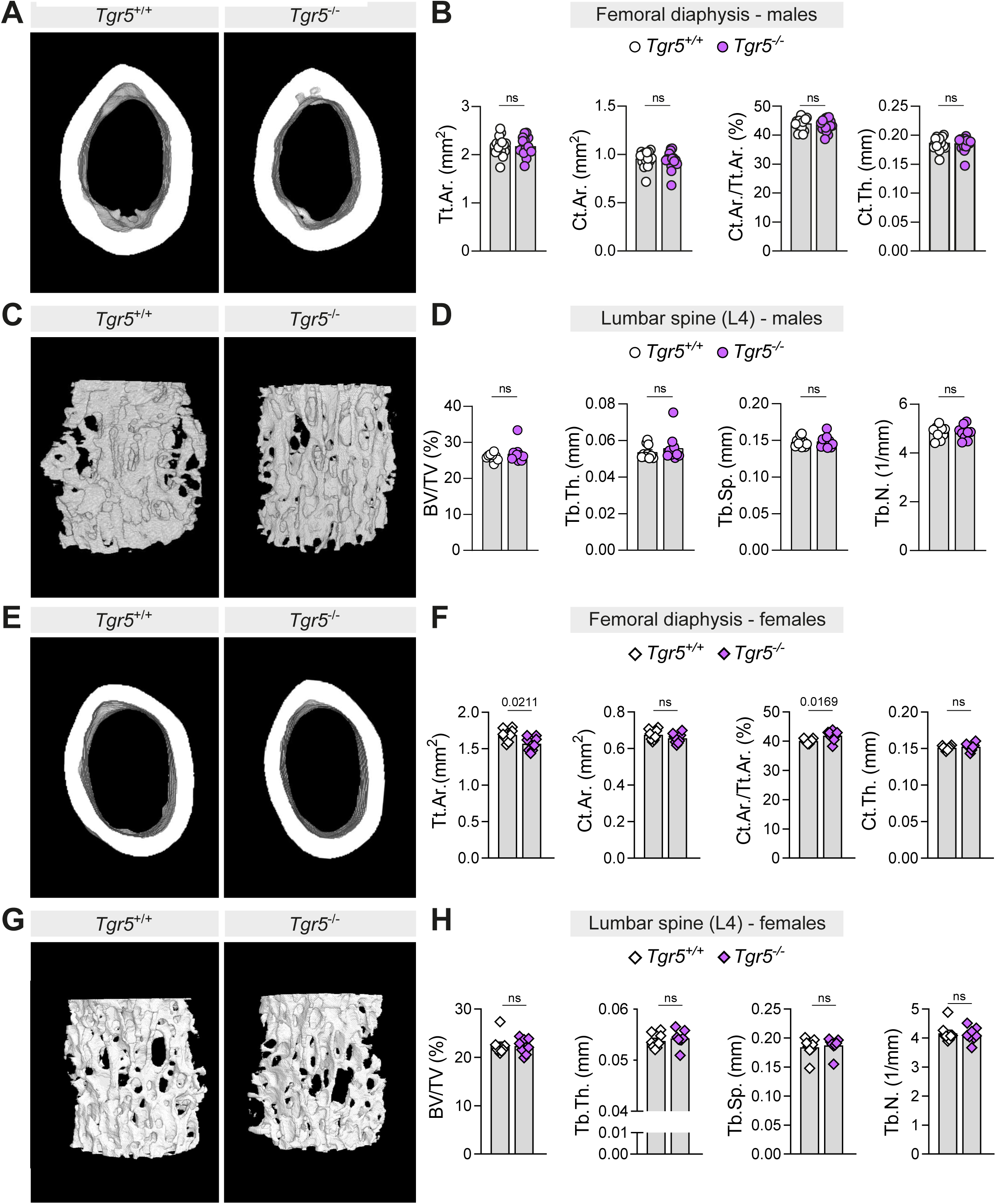
Loss of TGR5 did not alter the femoral cortical parameters or the spine microarchitecture. (**A**) Representative native µCT-derived 3D reconstructions of the cortical bone in the mid-femoral diaphysis of 8- to 12-week-old *Tgr5^+/+^* and *Tgr5*^−/−^ male mice. (**B**) µCT-derived bone morphometry measurements of the cortical bone in the mid-femoral diaphysis of 8- to 12-week-old *Tgr5^+/+^* and *Tgr5*^−/−^ male mice (n=17 for both *Tgr5^+/+^* and *Tgr5*^−/−^ mice); shown are the total cross-sectional area inside the periosteal envelope (Tt.Ar.), cortical bone area (Ct.Ar.), cortical area fraction (Ct.Ar./Tt.Ar.) and average cortical thickness (Ct.Th.). (**C**) Representative native µCT-derived 3D reconstructions of the trabecular structures in the fourth lumbar vertebrae (L4) of 8- to 12-week-old *Tgr5^+/+^* and *Tgr5*^−/−^ male mice. (**D**) µCT-derived bone morphometry measurements of the trabecular structures in L4 of 8- to 12-week-old *Tgr5^+/+^* and *Tgr5*^−/−^ male mice (n=10 for both *Tgr5^+/+^* and *Tgr5*^−/−^ mice); shown are bone volume/total volume (BV/TV), trabecular thickness (Tb.Th.), trabecular separation (Tb.Sp.) and trabecular number (Tb.N.). (**E**) Representative native µCT-derived 3D reconstructions of the cortical bone in the mid-femoral diaphysis of 8-week-old *Tgr5^+/+^* and *Tgr5*^−/−^ female mice. (**F**) µCT-derived bone morphometry measurements of the cortical bone in the mid-femoral diaphysis of 8-week-old *Tgr5^+/+^* and *Tgr5*^−/−^ female mice (n=8 for both *Tgr5^+/+^* and *Tgr5*^−/−^ mice); parameters shown analogously to **B**. (**G**) Representative native µCT-derived 3D reconstructions of the trabecular structures in the fourth lumbar vertebrae (L4) of 8-week-old *Tgr5^+/+^* and *Tgr5*^−/−^ female mice. (**H**) µCT-derived bone morphometry measurements of the trabecular structures in L4 of 8-week-old *Tgr5^+/+^* and *Tgr5*^−/−^ female mice (n=8 for both *Tgr5^+/+^* and *Tgr5*^−/−^ mice); parameters shown analogously to **D**. Results represent the mean ± s.e.m., n represents biologically independent replicates. Unpaired, two-tailed Student’s t-test **(B, D, F, H)** was used for statistical analysis. P values (exact value) are indicated, ns indicates no statistical significance.

**Figure 3-figure supplement 2.**
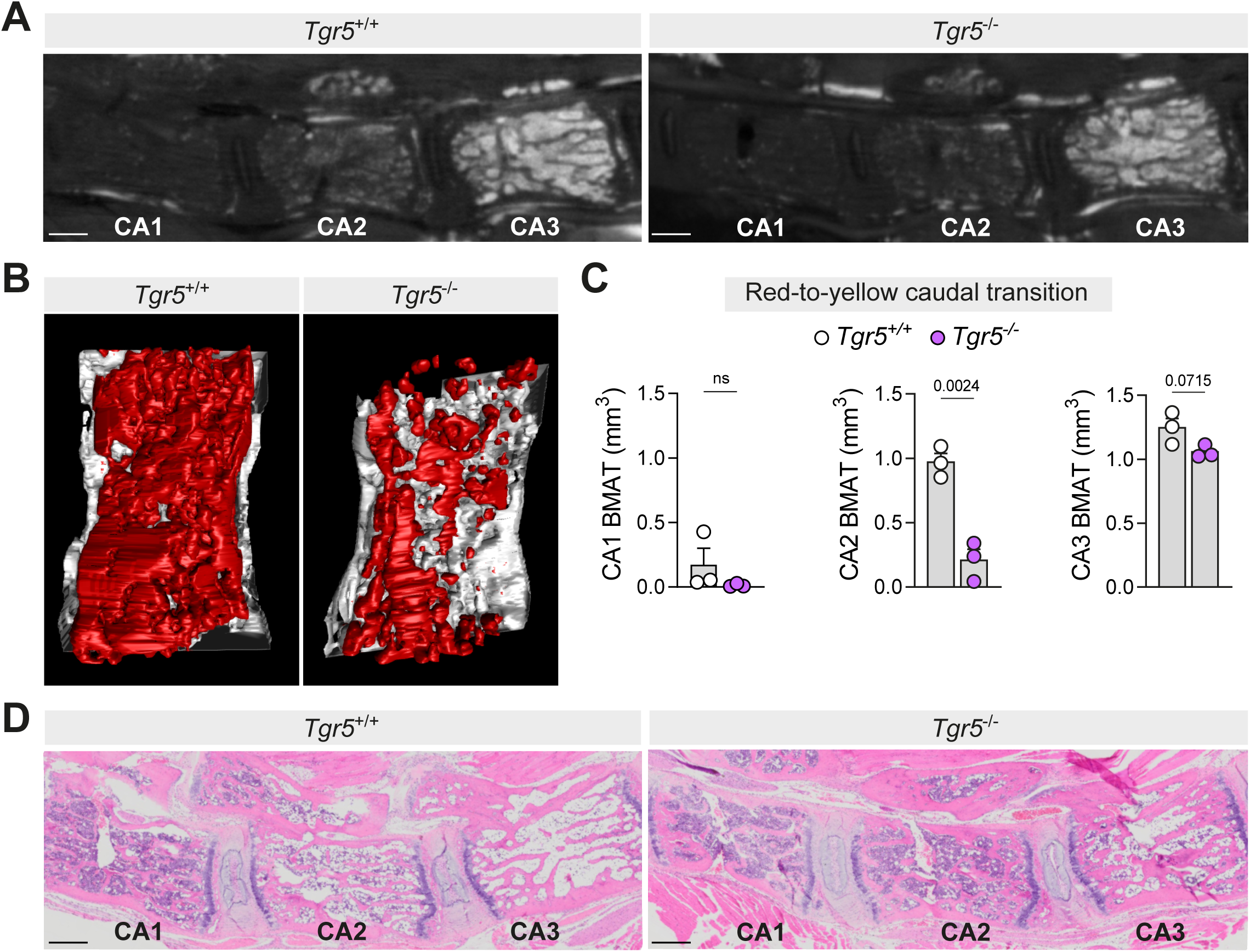
Loss of TGR5 alters the BM microenvironment in the caudal spine. (**A**) Representative images obtained from contrast-enhanced µCT scanning of decalcified OsO_4_-stained spines depicting lipid content in the red-to-yellow marrow transition in the caudal spine (CA) of 14-week-old male *Tgr5^+/+^* and *Tgr5*^−/−^ mice. (**B**) Sagittal view of a representative contrast-enhanced µCT-derived 3D reconstruction of OsO_4_-stained BMAT in the second caudal vertebrae (CA2) of 14-week-old *Tgr5^+/+^* and *Tgr5*^−/−^ male mice. (**C**) Quantification of the OsO_4_-stained BMAT for the first to third caudal vertebra (CA1-CA3) in the caudal red-to-yellow transition in 14-week-old *Tgr5^+/+^* and *Tgr5*^−/−^ male mice (n=3 for both *Tgr5^+/+^* and *Tgr5*^−/−^ mice). (**D**) Representative hematoxylin & eosin (H&E) histological sections of CA1-CA3 from a cohort independent from the animals shown in **A-C** (n=4 for both *Tgr5^+/+^* and *Tgr5*^−/−^ mice, 11- to 13-week-old). Scale bar in **A** and **D** = 500 µm. Results represent the mean ± s.e.m., n represents biologically independent replicates. Unpaired, two-tailed Student’s t-test **(C)** was used for statistical analysis. P values (exact value) are indicated, ns indicates no statistical significance.

**Figure 4-figure supplement 1.**
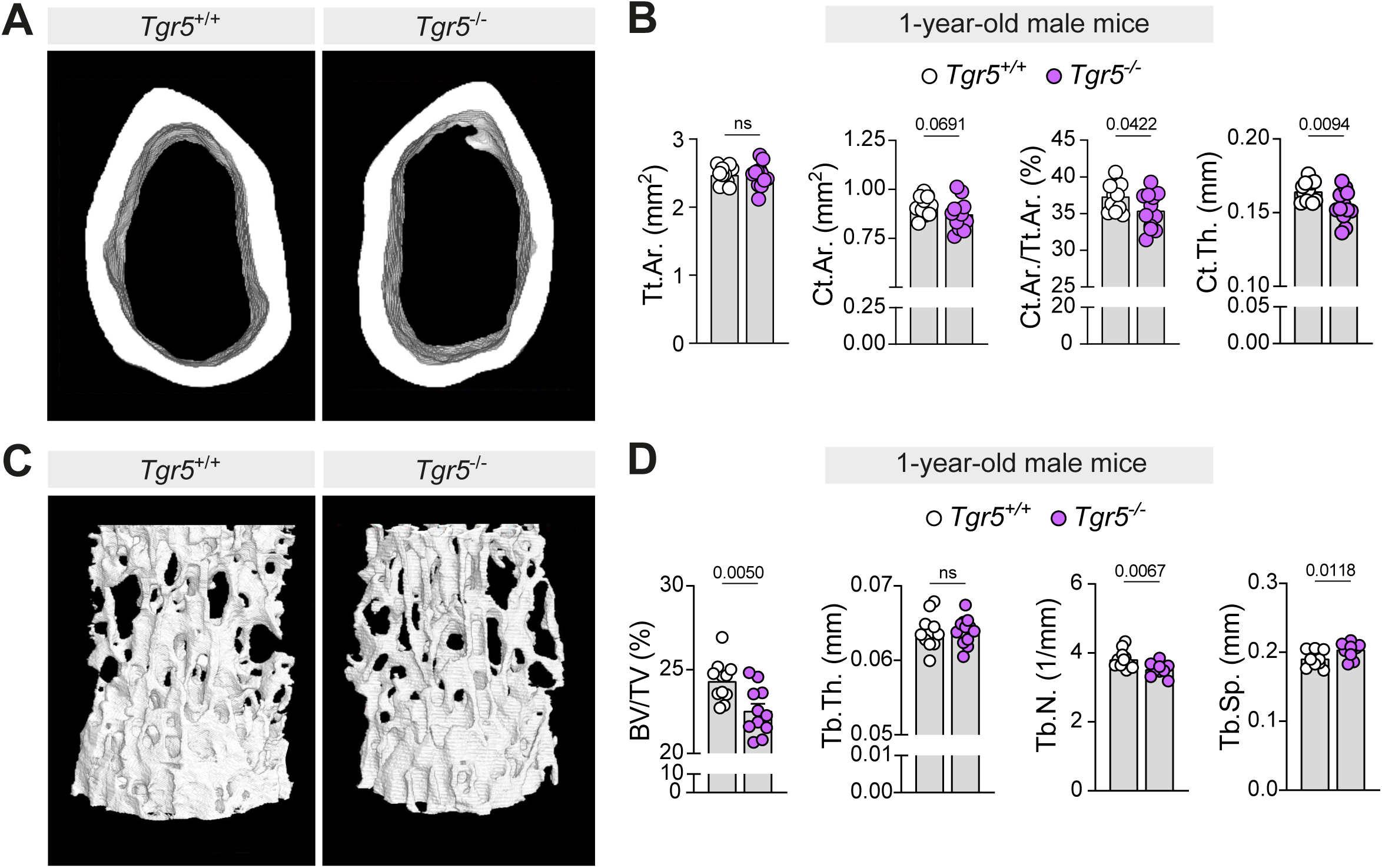
Loss of TGR5 slightly decreases vertebral and femoral cortical mineralized tissue in 1-year-old male mice. **(A)** Representative native µCT-derived 3D reconstructions of the cortical bone in the mid-femoral diaphysis of 1-year-old *Tgr5^+/+^* and *Tgr5*^−/−^ male mice. (**B**) µCT-derived bone morphometry measurements of the cortical bone in the mid-femoral diaphysis of 1-year-old *Tgr5^+/+^* and *Tgr5*^−/−^ male mice (n=12 for both *Tgr5^+/+^* and *Tgr5*^−/−^ mice); shown are the total cross-sectional area inside the periosteal envelope (Tt.Ar.), cortical bone area (Ct.Ar.), cortical area fraction (Ct.Ar./Tt.Ar.) and average cortical thickness (Ct.Th.). (**C**) Representative native µCT-derived 3D reconstructions of the trabecular structures in the fourth lumbar vertebrae (L4) of 1-year-old *Tgr5^+/+^* and *Tgr5*^−/−^ male mice. (**D**) µCT-derived bone morphometry measurements of the trabecular structures in L4 of 1-year-old *Tgr5^+/+^* and *Tgr5*^−/−^ male mice (n=12 for both *Tgr5^+/+^* and *Tgr5*^−/−^ mice); shown are bone volume/total volume (BV/TV), trabecular thickness (Tb.Th.), trabecular number (Tb.N.) and trabecular separation (Tb.Sp.). Results represent the mean ± s.e.m., n represents biologically independent replicates. Unpaired, two-tailed Student’s t-test (**B**, **D**) was used for statistical analysis. P values (exact value) are indicated, ns indicates no statistical significance.

**Figure 5-figure supplement 1.**
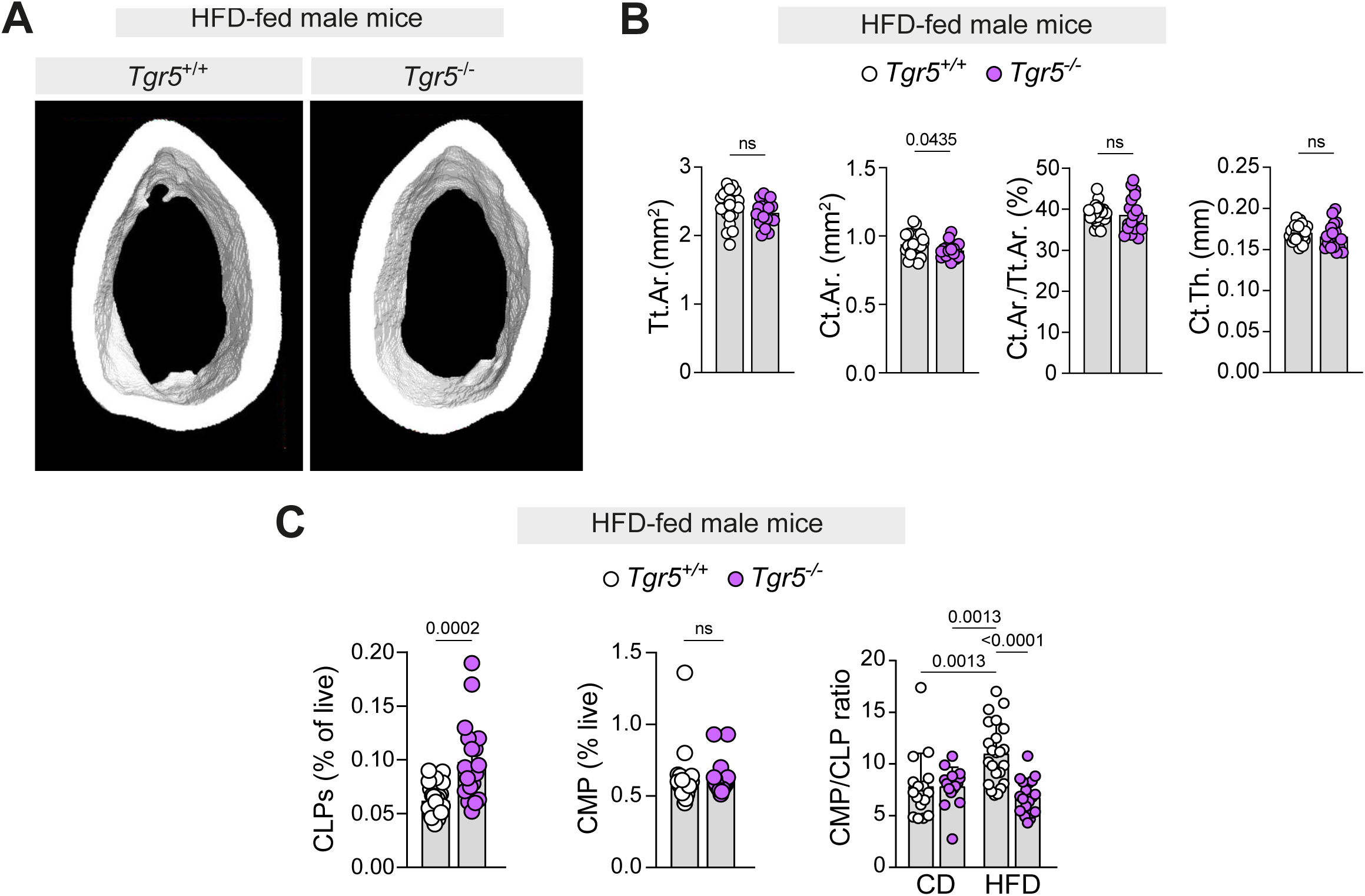
Loss of TGR5 does not affect femoral cortical parameters and blunts BM HSPC myeloid bias in HFD-fed male mice. (**A**) Representative native µCT-derived 3D reconstructions of the cortical bone in the mid-femoral diaphysis of HFD-fed (12 weeks) *Tgr5^+/+^* and *Tgr5*^−/−^ male mice. (**B**) µCT-derived bone morphometry measurements of the cortical bone in the mid-femoral diaphysis of HFD-fed (12 weeks) *Tgr5^+/+^* and *Tgr5*^−/−^ male mice (n=24 for *Tgr5^+/+^* and n=18 for *Tgr5*^−/−^ mice); shown are the total cross-sectional area inside the periosteal envelope (Tt.Ar.), cortical bone area (Ct.Ar.), cortical area fraction (Ct.Ar./Tt.Ar.) and average cortical thickness (Ct.Th.). (**C**) Frequency of common lymphoid progenitors (CLP), common myeloid progenitors (CMP) and ratio of CMP- to-CLP cells in the BM of CD-fed (calculated from the data presented in Figure 2A, same data as the “8- to 12-week-old” group in Figure 4F) and HFD-fed *Tgr5^+/+^* and *Tgr5*^−/−^ mice (n=24 for *Tgr5^+/+^* and n=20 for *Tgr5*^−/−^ mice). Results represent the mean ± s.e.m., n represents biologically independent replicates. Unpaired, two-tailed Student’s t-test was used for statistical analysis except for last panel in **C**, where a 2-way ANOVA test with Holm-Sidak’s multiple comparison correction was used. P values (exact value) are indicated, ns indicates no statistical significance.

**Figure 6-figure supplement 1.**
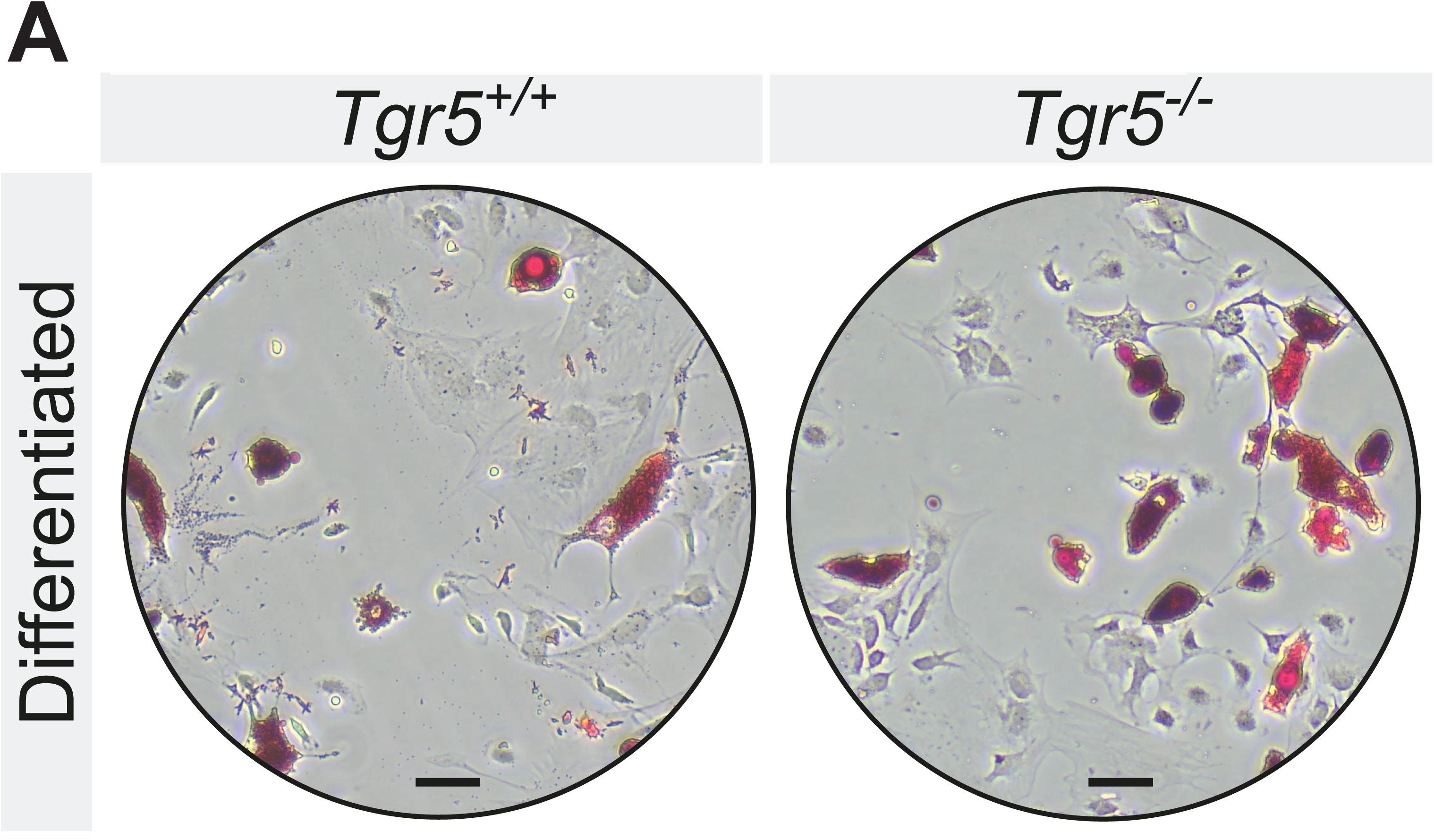
Loss of TGR5 increases in vitro BMSC differentiation into adipocytes. **(A)** Representative images of Oil red O (ORO)-stained BMSCs after 7 days of adipocytic differentiation (n=3 for both *Tgr5^+/+^* and *Tgr5*^−/−^ mice). Scale bar: 50 µm.

