## Supplementary Table 1 _ List of Antibodies for "The bile acid receptor TGR5 is a modulator of the bone marrow microenvironment"

| Antibodies flow cytometry (anti-mouse) | Commercial source | Cat.no |
|---|---|---|
| Biotin lineage cocktail | BD Biosciences | 51-9000724 |
| Streptavidin - Texas Red | Invitrogen | S872 |
| cKit - PE/Cy7 | Biolegend | 105814 |
| Sca1 - BV711 | BD Biosciences | 563992 |
| CD150 - PE/Cy5 | Biolegend | 115912 |
| CD48 - Pacific Blue | Biolegend | 103418 |
| CD34 - eFluor 660 | Invitrogen | 50-0314-82 |
| CD127 - PE | Invitrogen | 12-1271-83 |
| CD16/32 - Alexa Fluor700 | Invitrogen | 56-0161-82 |
| Sca1 - APC | Biolegend | 105112 |
| PDGFRa - PE | Biolegend | 135905 |
| CD45 - Alexa Fluor700 | Biolegend | 103128 |
| Ter119 - Alexa Fluor 700 | Biolegend | 116220 |
| CD24 APC eFluor780 | Invitrogen | 47-0242-82 |
| CD31 - PE/Cy7 | Biolegend | 102418 |
| CD45 - BV785 | Biolegend | 103149 |
| CD11b - BV650 | Biolegend | 101239 |
| Ly6G - BV711 | Biolegend | 127643 |
| Ly6C - Alexa Fluor 700 | Biolegend | 128024 |
| CD4 - PE/Cy5 | Biolegend | 100410 |
| CD19 - BV421 | BD Biosciences | 562440 |
| CD8 - BUV737 | BD Biosciences | 612759 |
| CD3 - APC/Cy7 | Biolegend | 100222 |
