## Supplementary material for "The bile acid receptor TGR5 is a modulator of the bone marrow microenvironment": Key Resources Table

| **Reagent type.** Duplicate rows as needed. Order is flexible, but row titles must be preserved. **Species** (mandatory if applicable). Append in parentheses. First mention, use full genus. **Sex** (if applicable). Add within parentheses after species. | **Designation.** Symbol/name used in publication. Separate multiple entries with semi-colon, space. (If semi-colon within designation, delimit wtith double quotes.) | **Source (public).** Stock center, company, data repository. **Reference.** Include PMID or DOI; use "this paper" if new. **If neither** applies enter "other" and explain in Additional Information. | **Identifiers.** Format as *ID_source:identifier* . Separate multiple entries with semi-colon, space. Include catalog numbers, stock numbers, database IDs or accession numbers, RRIDs. | **Additional information.** Mandatory if column C entry is "this paper" or "other". Description of new reagent; laboratory from which reagent obtained; other pertinent information. For antibodies, indicate dilution and conditions used. |
|---|---|---|---|---|

**Key Resources Table**

| Reagent type (species) or resource | Designation | Source or reference | Identifiers | Additional information |
|---|---|---|---|---|
| gene () | | NA | | |
| strain, strain background (*Mus musculus* ) | TGR5:GFP | | | |
| strain, strain background (*Mus musculus* ) | TGR5 knock-out mouse model (*Tgr5*$^{-/-}$ ) | | | |
| strain, strain background (*Mus musculus* ) | C57BL/6J (CD45.2) | | | |
| strain, strain background (*Mus musculus* ) | C57BL/6J Ly5.1 (CD45.1) | | | |
| strain, strain background (*Mus musculus* ) | congenic mice (CD45.1.2) | | | |
| genetic reagent () | | | | |
| cell line () | | | | |
| transfected construct () | | | | |
| biological sample () | | | | |
| antibody | Biotin lineage cocktail | BD Biosciences | 51-9000724 | |
| antibody | Streptavidin - Texas Red | Invitrogen | S872 | |
| antibody | cKit - PE/Cy7 | Biolegend | 105814 | |
| antibody | Sca1 - BV711 | BD Biosciences | 563992 | |
| antibody | CD150 - PE/Cy5 | Biolegend | 115912 | |
| antibody | CD48 - Pacific Blue | Biolegend | 103418 | |
| antibody | CD34 - eFluor 660 | Invitrogen | 50-0314-82 | |
| antibody | CD127 - PE | Invitrogen | 12-1271-83 | |
| antibody | CD16/32 - Alexa Fluor700 | Invitrogen | 56-0161-82 | |
| antibody | Sca1 - APC | Biolegend | 105112 | |
| antibody | PDGFRa - PE | Biolegend | 135905 | |
| antibody | CD45 - Alexa Fluor700 | Biolegend | 103128 | |
| antibody | Ter119 - Alexa Fluor 700 | Biolegend | 116220 | |
| antibody | CD24 APC eFluor780 | Invitrogen | 47-0242-82 | |
| antibody | CD31 - PE/Cy7 | Biolegend | 102418 | |
| antibody | CD45 - BV785 | Biolegend | 103149 | |
| antibody | CD11b - BV650 | Biolegend | 101239 | |
| antibody | Ly6G - BV711 | Biolegend | 127643 | |
| antibody | Ly6C - Alexa Fluor 700 | Biolegend | 128024 | |
| antibody | CD4 - PE/Cy5 | Biolegend | 100410 | |
| antibody | CD19 - BV421 | BD Biosciences | 562440 | |
| antibody | CD8 - BUV737 | BD Biosciences | 612759 | |
| antibody | CD3 - APC/Cy7 | Biolegend | 100222 | |
| antibody | GFP | Abcam | ab6673 | |
| antibody | | | | |
| recombinant DNA reagent | | | | |
| sequence-based reagent | | | | |
| peptide, recombinant protein | | | | |
| commercial assay or kit | SigmaFast | Sigma-Aldrich | | |
| commercial assay or kit | | | | |
| chemical compound, drug | Dafalgan | | | |
| chemical compound, drug | Osmium tetroxide ($OsO_4$) | | | |
| chemical compound, drug | Formalin | | | 10% |
| chemical compound, drug | EDTA | | | |
| chemical compound, drug | | | | |
| chemical compound, drug | | | | |
| chemical compound, drug | | | | |
| software, algorithm | Prism | GraphPad | Prism 9 | |
| software, algorithm | Custom CTan script | Bruker, Belgium | | |
| software, algorithm | NRecon | Bruker, Belgium | | |
| software, algorithm | Dragonfly software | ORS, Canada | | |
| software, algorithm | MATLAB | MathWorks, USA | | |
| other | Chow Diet | SAFE | SAFE 150 | |
| other | High Fat Diet | Research Diets | D12492 | |
| other | RS-2000 X-ray irradiator | RAD SOURCE | | |
| other | EDTA-coated tubes | | | |
| other | Element HT5 hematology analyzer | Heska, USA | | |
| other | SkyScanner 1276 | Bruker, Belgium | | |
| other | FACS buffer | | | 2% serum + 1 mM EDTA in PBS |
| other | PBS | | | |
| other | 3% $H_2O_2$ | | | |
| other | 0.1 M Tri-Na citrate | | | |
| other | ImmPRESS HRP | Vector Laboratories | | |
| other | 3,3'-Diaminobenzidine | Sigma-Aldrich | D5905 | |
| other | Harris hematoxylin | | | |
| other | Slide scanner | Olympus VS120-SL | | |
| other | Ascorbic-acid free αMEM | | | |
| other | FBS | | | |
| other | Penicillin-streptomycin | | | |
| other | Toluidine blue | | | |
| other | L-ascorbic acid | | | |
| other | β-glycerophosphate | | | |
| other | high-glucose (4.5 g/L) DMEM | | | |
| other | dexamethasone | | | |
| other | isobutylmethylxanthine | | | |
| other | insulin | | | |
| other | rosiglitazone | | | |
| other | Tween 20 | | | |
| other | alizarin red | | | |
| other | Oil Red O (ORO) | | | |
| other | iD3 plate reader | Agilent, USA | | |
| other | black wall 96-well imaging plates | Corning | 353962 | |
| other | Transmission DHM® T1000 | Lyncée Tec, Switzerland | | |
| other | Poly-ornithine | Sigma-Aldrich | | |
| other | | | | |
